# A generative network model reveals hierarchical structural memory guiding post-mitotic 3D genome inheritance

**DOI:** 10.64898/2026.10.28.762220

**Authors:** Leonardo Morelli, Stefano Cretti, Daniela Michelatti, Davide Bressan, Riccardo Scandino, Fulvio Chiacchiera, Davide Cittaro, Tiago P. Peixoto, Alessio Zippo

**Affiliations:** Department of Cellular, Computational and Integrative Biology (CIBIO), University of Trento, 38123 Trento, Italy; Center for Omics Sciences, IRCCS San Raffaele Scientific Institute, Milan, Italy; Center for Critical Computational Studies, Goethe University Frankfurt, 60629 Frankfurt am Main, Germany; Institute of Computer Science, Goethe University Frankfurt, 60629 Frankfurt am Main, Germany; IT:U Interdisciplinary Transformation University, 4040 Linz, Austria; Department of Network and Data Science, Central European University, 1100 Vienna, Austria; Complexity Science Hub, 1030 Vienna, Austria

## Abstract

How three-dimensional (3D) genome organization is reestablished after mitosis to preserve gene regulatory programs and cell identity remains unresolved. We asked whether mitotic chromosome architecture contributes to cellular memory by preserving structural information while maintaining the flexibility required for cellular adaptation. Here we introduce HiCONA, a generative Bayesian framework that infers hierarchical genome organization and quantifies the structural contribution of individual interactions through an Entropy Impact Score. HiCONA resolves genome organization across multiple scales, revealing structure-defining interaction patterns underlying chromatin architecture, independently of experimental modality. We identify structurally impactful chromatin interactions retained during mitosis delineating distinct trajectories of post-mitotic genome reassembly. These mitotically preserved interactions are enriched for active, transcriptionally competent regulatory elements, consistent with a cis-encoded architectural memory subsequently remodeled by trans-acting regulators. Together, our findings suggest that molecular information retained on mitotic chromosomes guides structural trajectories, providing an architectural basis for epigenetic memory and post-mitotic genome inheritance.

## Main

Epigenetic memory consists in the propagation of a transcriptional program throughout cell divisions, preserving cell identity beyond the instructive signals that originally established it^1^. The maintenance of such a transcriptional program results from the coordinated action of self-propagating molecular signatures encompassing both sequence-specific, *trans*-factors and *cis*-acting signals embedded within chromatin^2 3^. Cell division therefore represents a major challenge for properly preserving cell identity. During DNA replication, chromatin-associated information is diluted and must be accurately restored to preserve epigenetic information in daughter cells^4 5 6 7^. Mitosis poses an even greater challenge, as cells must not only ensure the faithful segregation of the genetic material but also preserve the epigenetic state while undergoing extensive reorganization of both two-dimensional chromatin composition and three-dimensional (3D) genome architecture^8^. Indeed, during mitosis most of the *trans*-factors including transcription factors and chromatin-associated regulatory proteins are either displaced from chromatin or degraded in response to mitotic signalling cascades. Likewise, many of the *cis*-acting signals associated with active chromatin state are transiently lost, whereas only a subset of genomic elements remains bookmarked, presumably to ensure their timely reactivation upon mitotic exit ^9 10 11 12 13 14 15^. In parallel, the 3D genome organization is extensively reshaped before being progressively reconstructed in daughter cells through mechanisms that remain only partially understood^16 17 18 19^. Current biophysical models^20^ and experimental data^21 22^ indicate that genome folding results from the combinatorial action of spatial compartmentalization and loop extrusion^19^. In the nuclear space chromatin is functionally segregated into A and B macro-compartments corresponding to transcriptionally active euchromatin and inactive heterochromatin regions, which are further organized into sub-compartments through preferential homotypic interactions among chromatin domains sharing similar epigenetic states^23 24^. Although histone modifications have been proposed to contribute to compartmentalization through mechanisms involving phase separation, the respective contributions of chromatin-intrinsic epigenetic information and trans-acting chromatin regulators in regulating homotypic interactions remain incompletely understood^25 26 27 28 29^. Nevertheless, the proposed contribution of histone modifications in genome folding implies that cell type-specific chromatin states delineate the 3D genome organization and how it gets transmitted to the progeny upon cell division^17 30 31 32 33 34^. In this context, loop extrusion participates to define and dynamically control the extent and frequency of contacts occurring among cis-regulatory elements (CREs) by constraining genome organization in topologically associated domains (TADs)^19 23 35^. Indeed, the cohesion-mediated loop extrusion in cooperation with structural proteins such as CTCFs delineate the space in which these contacts can occur, preserving cellular programs while ensuring responsiveness to environmental signals^36 37 38 39 40^. The functional relevance of CTCF-Cohesin-mediated genome folding is properly exemplified by mitotic transition during which these architectural components are displaced from chromatin, while Condensins determine a mitotic-specific genome folding program, which is pivotal to properly segregate the genomic material^19^. Upon mitotic exit, both CTCF and cohesion are progressively reloaded on chromatin to re-establish TADs boundaries and chromatin looping among CREs, while chromatin compartmentalization is properly restored only in the following G1 phase^17 18 30 31 32 33^. Recent studies have begun to dissect the earliest stages of post-mitotic genome reassembly, revealing previously unrecognized structural intermediates that emerge before the complete restoration of interphase genome architecture. Hansen and colleagues showed that transcriptionally competent CREs can form transient microcompartment-like interactions already during prometaphase that get strengthened in anaphase and telophase before being progressively remodeled during G1^30 31^. In parallel, Dekker and colleagues found that perturbation of nuclear import uncovers a chromatin-intrinsic folding program that precedes the activity of architectural and transcriptional machineries ^33^. Together, these observations show that a subset of chromatin interactions is preserved throughout mitosis, indicating that the early stages of post-mitotic genome refolding are characterized by chromatin-intrinsic structural intermediates that precede the complete restoration of interphase architecture. While these findings reveal an unexpected level of organization during the first stages of genome reassembly, their biological significance remains unclear. It is unknown whether these structural intermediates simply reflect transient configurations associated with chromosome reorganization or they actively contribute to transmitting the structural information required to faithfully reconstruct cell-type-specific genome architecture in daughter cells.

This uncertainty raises a fundamental conceptual paradox. Mitosis has traditionally been viewed as the greatest threat to epigenetic inheritance because it extensively reshapes both chromatin composition and three-dimensional genome organization ^2 18 19^. Yet every cell division faithfully reconstructs a cell-type-specific genome architecture despite the transient disappearance of many structural features. We therefore asked whether mitosis itself could actively contribute to cellular memory by preserving structural information while simultaneously maintaining sufficient configurational flexibility to permit changes in cell identity. We reasoned that this apparent contradiction is reminiscent of hysteretic physical systems^41 42^, in which the response of a system depends not only on its current molecular composition but also on the trajectory followed to reach its present state. Under this framework, post-mitotic genome reassembly would not simply result from *de novo* self-organization but would instead be guided by structural trajectories inherited through mitosis. Consequently, structural memory should not be encoded by the most abundant chromatin interactions but rather by spatial contacts contributing most strongly to maintaining the stability of the folding landscape.

Addressing this hypothesis requires moving beyond the current interpretation of chromosome conformation data. Although remarkable advances in chromosome conformation capture and imaging technologies have enabled genome organization to be resolved across multiple spatial scales, most computational approaches remain inherently static^43 44 45^. They identify loops, TADs, compartments and microcompartments as independent architectural features but do not distinguish interactions that merely occur from those that preserve the structural organization of the genome during dynamic transitions such as mitotic exit^43 44 45 46^. Consequently, current analytical frameworks provide limited insight into how structural information evolves throughout the cell cycle, which interactions contribute to genome inheritance and how previous chromatin configurations influence subsequent folding trajectories ^47 46^.

To overcome these limitations, we developed HiCONA (Hi-C Organization with Network Analysis), a generative Bayesian framework that reconstructs genome architecture as a probabilistic structural hierarchy and quantifies the structural contribution of individual chromatin interactions through the Entropy Impact Score (EIS) ^48 49^. Rather than describing genome organization as a collection of independent architectural entities, HiCONA identifies the interactions that contribute most strongly to minimizing the structural entropy of chromatin organization, thereby providing a quantitative framework to identify the structural trajectories that govern post-mitotic genome reassembly.

## Results

### Probabilistic reconstruction identifies structure-defining chromatin interactions

To identify the interactions that encode structural information within chromosome conformation maps, we developed HiCONA (Hi-C Organization with Network Analysis). Starting from an imaging- or sequencing-based chromatin contact maps, HiCONA infers the hierarchical chromatin organization most consistent with the observed interaction patterns and subsequently quantifies the contribution of every chromatin interaction to this organization through the Entropy Impact Score (EIS) (Fig. 1a). Unlike contact frequency, which measures how often two loci interact, EIS measures how strongly individual interactions constrain the inferred genome organization, thereby providing a complementary view of chromosome architecture.

**Fig. 1.**
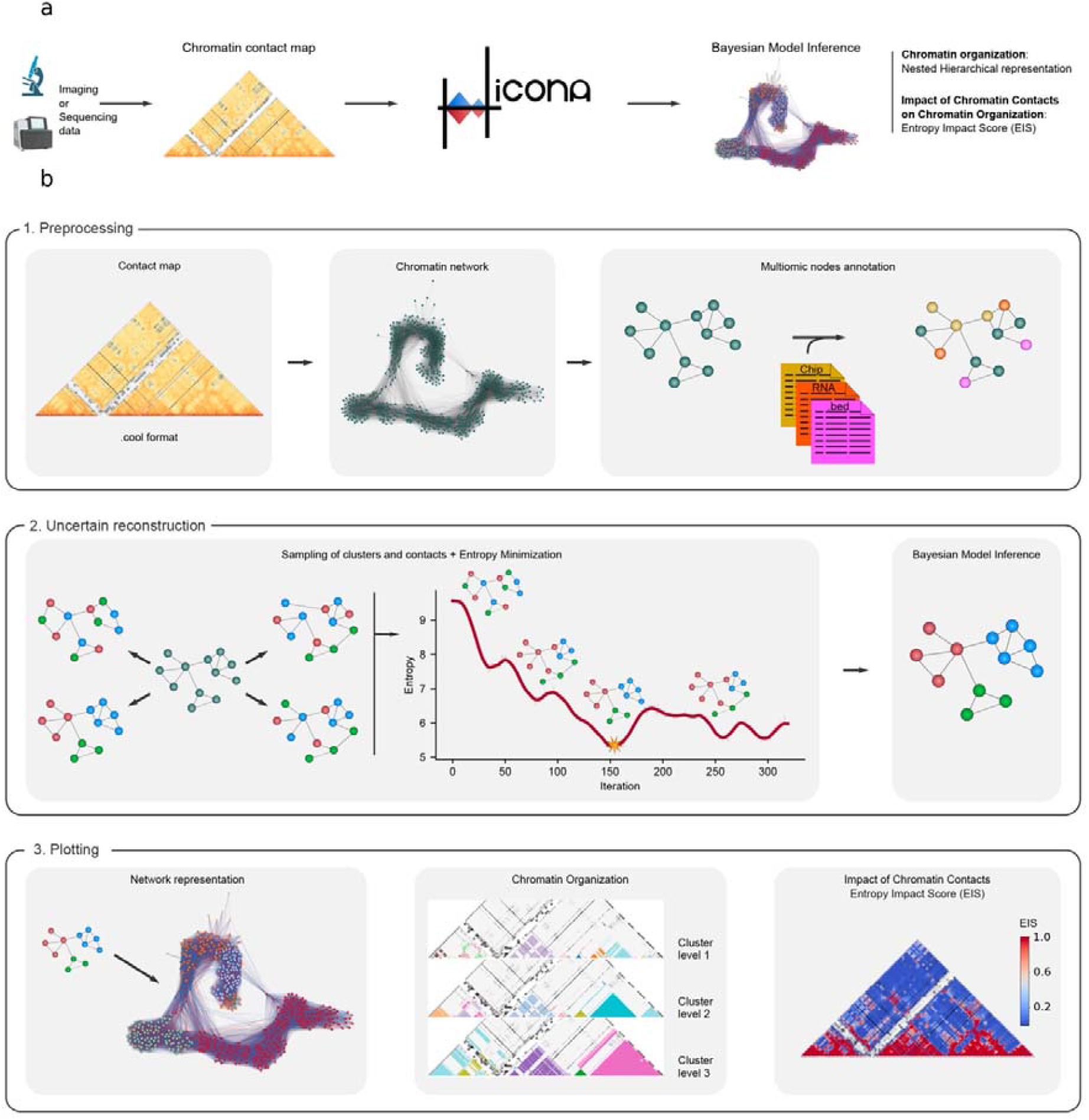
HiCONA identifies structure-defining chromatin contacts through probabilistic reconstruction of genome architecture a. Schematic workflow of HiCONA. Starting from chromatin contact maps generated by either imaging or sequencing based approach, HiCONA reconstructs nested hierarchical chromatin organization and quantifies the contribution of individual contacts to genome architecture through the Entropy Impact Score (EIS). **b**. In the preprocessing step, contact maps in .cool format are converted into chromatin networks, in which nodes represent genomic loci and edges represent chromatin contacts; nodes can be further annotated with multi-omic features. During uncertain reconstruction, alternative network configurations are explored by stochastic sampling, and entropy minimization is used to identify the simplest organization among models with the highest explanatory power. Bayesian model inference generates a probabilistic representation of chromatin architecture that captures uncertainty in cluster assignments and contact patterns. The inferred network is then used to derive hierarchical chromatin organization and to calculate EIS, which measures the impact of individual chromatin contacts on the overall organizational entropy of the system. Results can be visualized as network representations, hierarchical contact maps, or contact-impact heatmaps.

HiCONA first represents chromosome conformation as a weighted interaction network in which genomic loci constitute vertices and chromatin contacts define weighted edges (Fig. 1b, top). Functional genomic features, including epigenomic and transcriptional annotations, can be incorporated as node attributes, allowing chromosome organization to be interpreted within its biological context.

The hierarchical organization of the chromatin network is inferred using a degree-corrected nested stochastic block model (nSBM), a Bayesian generative model that recursively partitions the interaction network into nested communities by minimizing its description length ^48 49^ (Fig. 1b, center). Rather than identifying individual architectural features such as loops, topologically associating domains or compartments independently, the model infers the latent hierarchical organization that best explains the observed interaction patterns, allowing these structural features to emerge naturally from the inferred organization without requiring predefined assumptions^48 49^. Because multiple hierarchical organizations can explain the same chromosome conformation map with comparable statistical support, HiCONA samples the posterior distribution of genome organizations instead of relying on a single optimal solution^48 49^. Using Markov chain Monte Carlo inference, the algorithm^50^ explores alternative hierarchical reconstructions while preserving their relative posterior probabilities, generating an ensemble of plausible chromosome organizations consistent with the observed data. This probabilistic formulation explicitly captures the uncertainty inherent to chromosome conformation measurements while distinguishing organizational features that remain robust across alternative reconstructions ^51^.

We reasoned that chromatin interactions consistently required across this posterior ensemble should represent fundamental organizational constraints, whereas interactions compatible with many alternative reconstructions should contribute comparatively little to maintaining genome architecture. Based on this principle, we developed the EIS (Fig. 1b, center), which quantifies the structural contribution of every chromatin interaction according to its influence on the posterior ensemble of inferred genome organizations. Consequently, structurally informative interactions are not necessarily the most abundant contacts detected by chromosome conformation assays, but those whose removal most strongly alters the inferred organizational landscape. Finally, HiCONA summarizes these probabilistic reconstructions through complementary representations, including hierarchical chromatin networks, multiscale structural partitions and genome-wide maps of EIS values (Fig. 1b, bottom). HiCONA’s hierarchy comprises fine clusters at the lowest level, nested within progressively fewer, larger clusters at higher levels. By transforming chromosome conformation maps into probabilistic structural landscapes, HiCONA enables chromatin interactions to be evaluated according to their contribution to genome organization rather than their interaction frequency. We next asked whether these structurally informative interactions recapitulate established principles of chromosome folding and reveal organizational features that remain inaccessible using conventional chromosome conformation analysis.

### Hierarchical chromatin organization captures distinct physical principles of genome folding

To determine whether the hierarchy inferred by HiCONA reflects established mechanisms of chromosome folding, we first generated polymer models in which loop extrusion and compartmentalization could be independently modulated^52^. We simulated 4,000 monomers in each polymer model, corresponding to approximately 1017Mb of genomic sequence, and assigned each monomer to either the A or B compartment. Compartmentalization was varied by increasing the strength of homotypic attraction between monomers belonging to the same compartment, whereas loop extrusion was tuned by varying the release probability of designated loop extruders, with lower release probabilities producing more stable extrusion-driven domains (Fig. 2a, b and Extended Data Fig. 1).

**Fig. 2.**
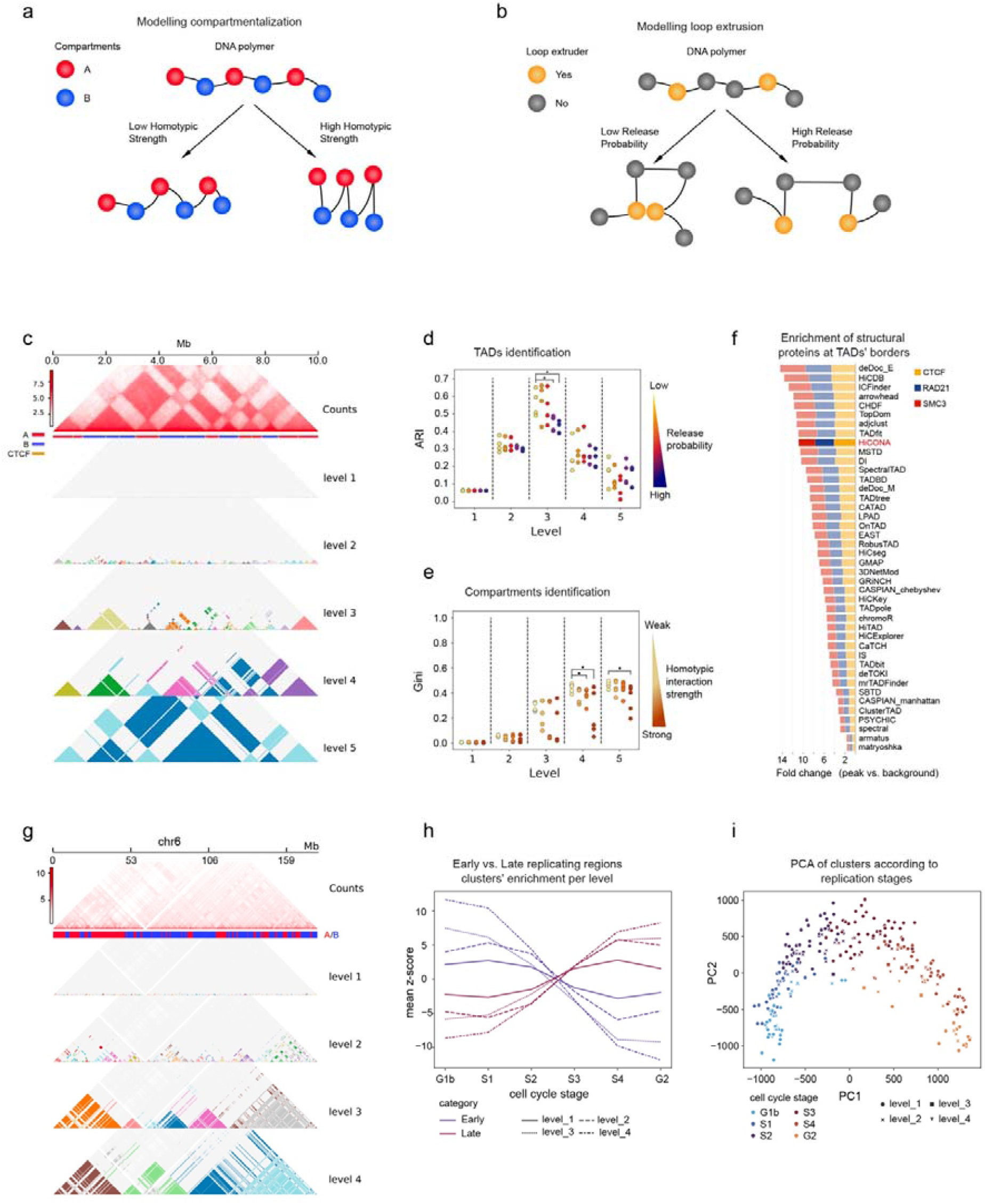
HiCONA recovers hierarchical chromatin organization in simulated and experimental data a. Schematic of compartmentalization simulations comprising A (red) and B (blue) chromatin compartments. Representative simulated contact maps generated with low homotypic strength (HS; left) and high HS (right) are shown below. Schematic of loop-extrusion simulations involving extruder molecules (yellow). **b**. Representative simulated contact maps generated with high release probability (RP) of two extruders (left) and low RP (right) are shown below. **c**. Simulated chromatin polymer model consisting of 4,000 monomers representing 10 Mb of chromatin, generated with RP = 1.0 and HS = 3. Top, simulated contact map (log2 counts). Middle, linear annotation of compartment identities and loop-extruder positions. Bottom, nested hierarchical chromatin organization inferred by HiCONA. **d**. Adjusted Rand index (ARI) between cluster boundaries identified at each hierarchical level and simulated loop-extruder positions across combinations of RP and HS parameters. **e**. Gini index measuring compartmental homogeneity of clusters at each hierarchical level across combinations of RP and HS parameters. **f**. Enrichment of CTCF, RAD21, and SMC3 occupancy at topologically associating domain (TAD) boundaries identified by HiCONA and 39 alternative TAD-calling methods on chromosome 6 of GM12878 cells. **g**. HiCONA hierarchy, inferred from chromosome 6 of GM12878 cells at 50-kb resolution (log2 counts), illustrating concordance between hierarchical organization and chromatin compartment structure. **h**. Mean Repli-seq signal for clusters identified at each hierarchical level, comparing early- and late-replicating domains. **i**. Principal component analysis (PCA) of all clusters across hierarchical levels of chromosome 6 in GM12878 cells based on enrichment for Repli-seq signals.

HiCONA reconstructed a nested hierarchy from each simulated contact map, allowing us to determine whether the structural features generated by these two mechanisms emerged at distinct organizational levels (Fig. 2c). To determine which level of the inferred hierarchy best recapitulated the simulated loop-extrusion domains, we quantified partition similarity using the adjusted Rand index (ARI). The third hierarchical level showed the strongest correspondence, with agreement increasing progressively as extruder release probability decreased (Fig. 2d). To assess compartment recovery, we quantified the homogeneity of A and B labels within each cluster using the Gini index. Small clusters at lower hierarchical levels were compartmentally homogeneous across conditions. However, stronger homotypic attraction extended this homogeneity to coarser levels of the hierarchy, with the most pronounced effects at levels 4 and 5. (Fig. 2e). The complete parameter space produced consistent hierarchical transitions, with extrusion-driven domains emerging at intermediate levels and large-scale compartmental organization becoming detectable at upper levels only when homotypic attraction was sufficiently strong (Extended Data Fig. 1). Notably, when applied to a larger polymer model representing a 30-Mb chromatin region (Extended Data Fig. 2a), HiCONA correctly identifies TADs at the second level of the hierarchy (Extended Data Fig. 2b), while maintaining a clear separation between compartments across hierarchical levels (Extended Data Fig. 2c). Thus, HiCONA resolved the relative contributions of two distinct physical mechanisms and localized their structural effects to different levels of the same hierarchy.

We next asked whether the latent hierarchy inferred by HiCONA reflects the organizational scales of chromosome folding depicted by experimental approaches. Using Hi-C data from chromosome 6 in GM12878 cells, we benchmarked the lowest hierarchical level against 39 dedicated TAD-calling approaches through TADshop^53^ (Fig. 2f and Extended Data Fig. 2d, e). Despite not being optimized specifically for TAD detection, HiCONA ranked 13th overall across orthogonal measures of domain quality, including boundary insulation, within-domain contact enrichment, histone-modification segregation and enrichment of architectural proteins at domain boundaries (Fig. 2f and Extended Data Fig. 2d). HiCONA effectively recovered boundaries enriched for CTCF, RAD21 and SMC3, ranking ninth among the evaluated approaches (Fig. 2f), and identified domains with significant H3K27me3/H3K36me3 segregation (Extended Data Fig. 2d). Thus, the lowest level of the HiCONA hierarchy recapitulates conventional mesoscale organization without feature-specific definitions or a dedicated TAD-calling procedure.

We next asked whether higher levels of the HiCONA hierarchy capture compartment-scale organization across chromosomes 6, 13 and 21, which differ substantially in genomic length (Fig. 2g and Extended Data Fig. 2f-g). We first characterized the inferred clusters using replication timing, a functional correlate of A/B compartment identity^54^. Principal-component analysis of Repli-seq profiles separated clusters according to their representative replication stage, while the mean profiles became progressively polarized toward early- or late-replicating chromatin at higher hierarchical levels (Fig. 2h, i and Extended Data Fig. 3h–k). These results indicate that broader levels of the HiCONA hierarchy resolve functionally distinct chromatin environments associated with active and repressive compartments. We then used the Gini index to quantify the extent to which individual clusters were dominated by a single A/B compartment identity. At lower and intermediate hierarchical levels, 90–95% of loci within each cluster were typically assigned to the same compartment, corresponding to Gini values of 0.10–0.18. This correspondence remained evident at the upper levels of chromosomes 6 and 13, where Gini values of approximately 0.32 indicated that most loci still shared the same compartment assignment, although less consistently than across the hierarchy of chromosome 21 (Extended Data Fig. 3l). This difference is consistent with the larger genomic spans encompassed by the upper hierarchical levels of chromosomes 6 and 13. Together, the replication-timing profiles and distributions of A/B identities show that the upper levels of the HiCONA hierarchy capture large-scale, functionally coherent chromatin organization across chromosomes of different sizes. The hierarchy therefore captured functionally distinct large-scale chromatin environments in addition to local topological domains. Notably, the strongest correspondence with TADs occurred at level 3 in the 2.5-kb-resolution polymer simulations but at level 1 in 10-kb experimental Hi-C data, consistent with finer-resolution simulations resolving sub-TAD structures below complete domains. Together, these analyses establish that the latent hierarchy inferred by HiCONA is not an arbitrary network partition but a multiscale representation of chromosome architecture. Lower levels recovered extrusion-driven and mesoscale domains, whereas higher levels captured compartmental segregation and its functional association with replication timing. The correspondence between hierarchical level and structural scale depended on input resolution, with the finer polymer simulations resolving sub-TAD structures below complete extrusion domains. These results therefore validate the physical and biological organization on which EIS is computed. We next directly tested whether EIS distinguishes interactions that constrain this organization from contacts that are frequent but structurally dispensable.

**Fig. 3.**
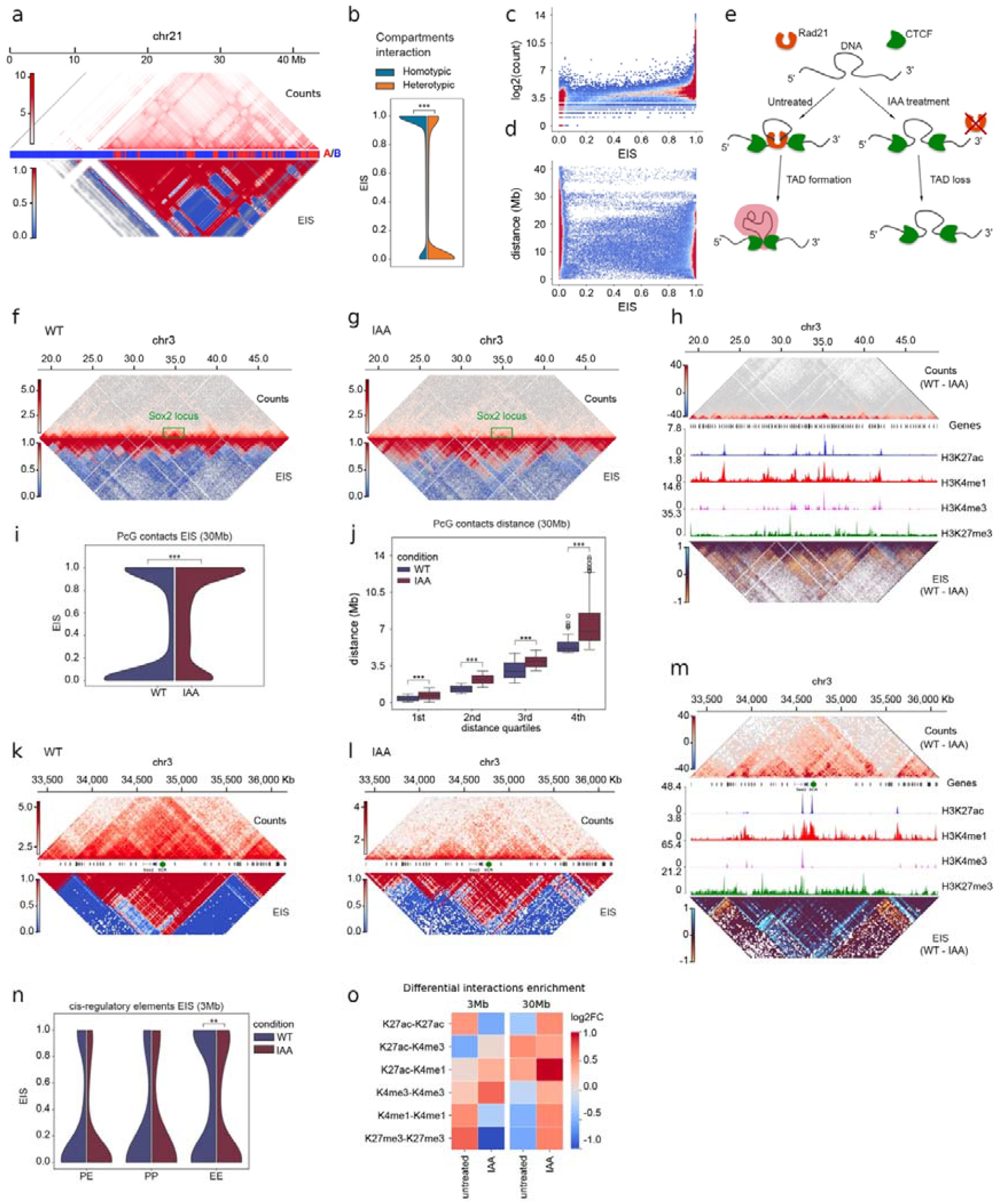
Entropy Impact Score identifies structure-defining chromatin contacts and reveals their reorganization following cohesin depletion a. Chromosome 21 of GM12878 contact log2 counts (top) and EIS (bottom) resolution 50kb. **b**. EIS of homotypic and heterotypic compartment interaction in chromosome 21 of GM12878 cells computed at 50kb. **c**. Density plot showing the distribution of chromatin contacts as a function of EIS and the number of observed contacts (log2-transformed counts). Higher densities are indicated by warmer colors. **d**. Density plot showing the distribution of chromatin contacts as a function of EIS and genomic separation. **e**. Schematic representation of chromatin organization in the presence and absence of cohesin. **f**. chr3:18,000,000–49,000,000 in mESC cells log2 contact counts (top) and EIS (bottom) in untreated cells at 40kb resolution. **g**. chr3:18,000,000–49,000,000 in mESC cells log2 contact counts (top) and EIS (bottom) in IAA-treated cells at 40kb resolution. **h**. Differential contact maps of untreated vs IAA mESC cells (chr3:18,000,000–49,000,000) using counts (top) or EIS (bottom). Genes and normalized read counts of H3K27ac, H3K4me1, H3K4me3 and H3K27me3 are shown. **i**. EIS distribution of H3K27me3-H3K27me3 contacts in untreated and IAA conditions in chr3:18,000,000–49,000,000 of mESC cells (P-value computed with two-sided Mann-Whitney U test). **j**. Contact distance of H3K27me3-H3K27me3 high-impact interactions (EIS>0.5) in chr3:18,000,000–49,000,000 of mESC cells in untreated and IAA conditions, divided by quartile distances. Cliff’s delta effect size: 1st quartile = 0.40, 2nd quartile = 0.89, 3rd quartile = 0.57, 4th quartile = 0.79. P-values were computed with Mann–Whitney U test. **k**. Sox2 locus (chr3:33,390,000–36,180,000) in mESC cells log2 contact counts (top) and EIS (bottom) in untreated cells at 20kb resolution. Sox2 Promoter and SCR are shown (center). **l**. Sox2 locus (chr3:33,390,000–36,180,000) in mESC cells log2 contact counts (top) and EIS (bottom) in IAA-treated cells at 20kb resolution. Sox2 Promoter and SCR are shown (center). **m**. Differential contact maps of untreated vs IAA mESC cells (chr3:33,390,000–36,180,000) using counts (top) or EIS (bottom). Genes and normalized read counts of H3K27ac, H3K4me1, H3K4me3 and H3K27me3 are shown. **n**. EIS of Promoter-Promoter (left), Promoter-Enhancer (center), and Enhancer-Enhancer (EE) interactions in untreated cells and IAA cells (p-value computed with two-sided Mann-Whitney U test). **o**. Heatmap showing the log2 fold change (log2FC) in interaction enrichment between genomic regions carrying distinct histone modifications at 3 Mb and 30 Mb genomic scales in untreated and IAA-treated cells. Rows indicate pairs of chromatin states defined by H3K27ac, H3K4me3, H3K4me1, and H3K27me3 marks. Positive values (red) indicate enriched interactions, whereas negative values (blue) indicate depleted interactions relative to the expected interaction frequency.

### Structure-defining chromatin interactions uncover distinct principles of genome organization beyond contact frequency

Having established that the HiCONA hierarchy recapitulates the multiscale organization of chromosomes, we next asked whether EIS identifies individual interactions with established roles in maintaining chromosome architecture. We first computed chromosome-scale EIS maps for chromosome 21 in GM12878 cells and compared homotypic interactions connecting loci belonging to the same A/B compartment with respect to heterotypic interactions spanning opposite compartments (Fig. 3a). Homotypic interactions exhibited significantly higher EIS values than heterotypic ones, with 74–79% of homotypic contacts assigned EIS values between 0.8 and 1 across maps generated at 50-kb, 100-kb and 250-kb resolution (Fig. 3b and Extended Data Fig. 3a–c). This enrichment was reproduced in IMR90 cells across the same resolution range, indicating that the preferential assignment of high EIS to homotypic compartment interactions was independent from the cellular model and data resolution (Extended Data Fig. 3d–g). Importantly, high-EIS interactions occurred across a broad range of contact frequencies and genomic distances, showing that EIS is not a simple transformation of either contact abundance or linear separation (Fig. 3c, d and Extended Data Fig. 3h).

We next exploited time-controlled RAD21 depletion through degron-based modality^55^ as a defined perturbation that weakens cohesin-dependent loop extrusion while reinforcing compartmental segregation (Fig. 3e). Across a 30-Mb region of chromosome 3, untreated mESCs displayed triangular domains of high EIS, mirroring the characteristic TAD-based organization (Fig. 3f). Following RAD21 depletion, this pattern was replaced by extended and long-range blocks of structurally impactful interactions, consistent with a redistribution of structural importance from local TAD-based organization toward compartment-associated folding.^21^ (Fig. 3g). Differential contact frequencies predominantly emphasized the loss of short-range interactions within domains, with comparatively few contacts gaining abundance after depletion. By contrast, differential EIS analyses revealed a broader redistribution of structural importance: contacts preferentially contributing to the 3D organization in the wild type mESCs were predominantly short-range interactions, whereas those gaining impact after RAD21 depletion extended over larger genomic distances and frequently crossed TADs boundaries (Fig. 3h). The corresponding HiCONA hierarchies recapitulated this transition, with TAD-like higher-level domains in untreated cells being replaced by broader, compartment-like partitions after cohesin loss (Extended Data Fig. 4a, b).

**Fig. 4.**
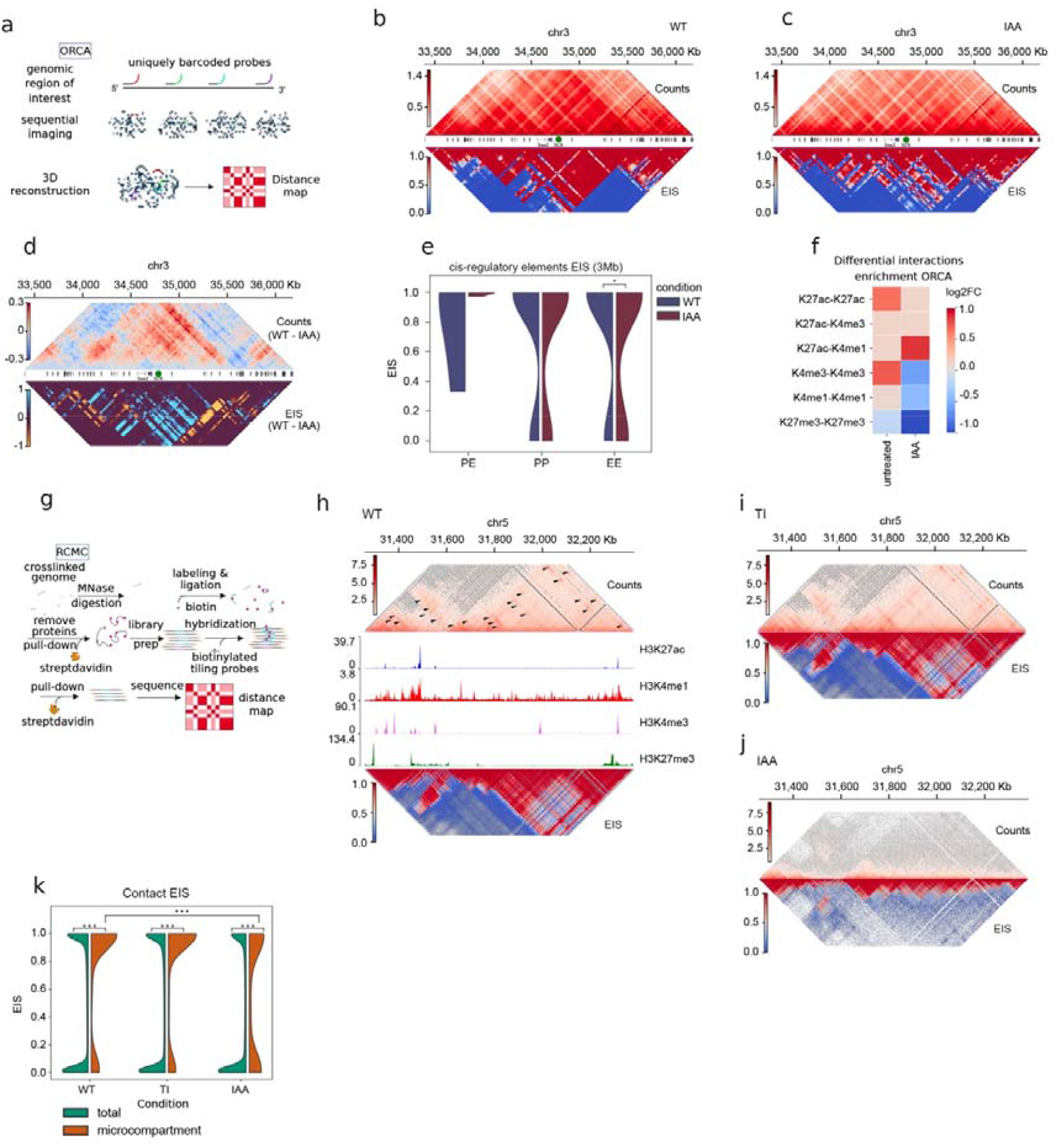
Entropy Impact Score identifies structure-defining chromatin interactions across experimental modalities and genomic scales a. Schematic representation of ORCA imaging technique. **b**. Log2 contact counts (top) and EIS (bottom) for the Sox2 locus (chr3:33,390,000–36,180,000) in mESCs, shown for untreated cells (ORCA). This region was probed using 74 probes, yielding an approximate resolution of ~37 kb. **c**. Log2 contact counts (top) and EIS (bottom) for the Sox2 locus (chr3:33,390,000–36,180,000) in mESCs, shown for IAA-treated cells (ORCA). This region was probed using 74 probes, yielding an approximate resolution of ~37 kb. **d**. Differential contact maps of untreated vs IAA mESC cells (chr3:33,390,000–36,180,000) using counts (top) or EIS (bottom). **e**. ORCA EIS of Promoter-Enhancer (left), Promoter-Promoter (center), and Enhancer-Enhancer (EE) interactions in untreated cells and IAA cells (p-value computed with two-sided Mann-Whitney U test). **f**. Heatmap showing the log2 fold change (log2FC) in interaction enrichment between genomic regions carrying distinct histone modifications in untreated and IAA-treated cells. Rows indicate pairs of chromatin states defined by H3K27ac, H3K4me3, H3K4me1, and H3K27me3 marks. Positive values (red) indicate enriched interactions, whereas negative values (blue) indicate depleted interactions relative to the expected interaction frequency. **g**. Schematic representation of RCMC sequencing technique. **h**. chr5:31300000-32400000 of mESC cells log2 contact counts (top) and EIS (bottom) in untreated cells at 1kb resolution (arrows represent some of the microcompartments). Normalized read counts of H3K27ac, H3K4me1, H3K4me3 and H3K27me3 are shown. **i**. chr5:31300000-32400000 of mESC cells log2 contact counts (top) and EIS (bottom) in transcription-inhibited cells at 1kb resolution. **j**. chr5:31300000-32400000 of mESC cells log2 contact counts (top) and EIS (bottom) in cohesin depleted cells at 1kb resolution. **k**. EIS of all contacts and microcompartment interactions in chr5:31300000-32400000 of mESC cells at 1kb in untreated, TI and IAA conditions (P-value computed with two-sided Mann-Whitney U test).

Because interactions between Polycomb-associated regions can support long-range folding independently of loop extrusion^55^, we next examined homotypic contacts between H3K27me3-enriched regions. These interactions showed a global increase in EIS following RAD21 depletion (Fig. 3i). Among high-impact contacts, H3K27me3-associated interactions also spanned significantly greater genomic distances in depleted cells, with the strongest separation observed among contacts in the most distal group (Fig. 3j). Thus, when local cohesin-dependent organization was impaired, long-range contacts between Polycomb (PcG) chromatin domains acquired greater relative structural importance.

Having identified a redistribution of structural importance from local domain organization toward long-range compartment- and Polycomb-associated interactions, we next asked how this transition affects the structural contribution of cis-regulatory interactions within a defined regulatory domain. We focused on the Sox2 locus, a well-characterized regulatory domain containing multiple cis-regulatory elements, including the distal Sox2 Control Region (SCR). Acute cohesin depletion strongly disrupts TAD-scale organization and reduces long-range regulatory contacts, including Sox2–SCR interactions, without completely abolishing CRE communication within the locus ^56 57^. In untreated cells, the EIS landscape delineated the Sox2 TAD as a coherent domain of high-impact interactions (Fig. 3k). RAD21 depletion weakened this organization and redistributed structural impact toward contacts extending beyond the original TAD boundaries (Fig. 3l). Consistently, the Sox2 domain was recovered at multiple levels of the untreated HiCONA hierarchy but was no longer detectable following depletion (Extended Data Fig. 4c, d). Differential contact frequencies primarily captured the loss of interactions within the Sox2 TAD, whereas differential EIS additionally identified contacts that gained structural importance across its former boundaries (Fig. 3m). EIS therefore revealed not only the erosion of local insulation but also the alternative interactions acquiring structural relevance following cohesin loss. Of note, a comparable reorganization was observed across an independent region of chromosome 6 (Extended Data Fig. 4e-j). Given that regulatory interactions within the Sox2 locus are reduced but not completely abolished following cohesin depletion, we next asked whether CRE contacts that remain detectable retain their structural contribution within the reorganized chromatin network. Enhancer–enhancer interactions showed significantly lower EIS after RAD21 depletion, while both promoter–enhancer and promoter-promoter contacts remained largely stable across both conditions (Fig. 3n). Chromatin-state annotation of differential EIS contacts further supported a scale-dependent redistribution in which local regulatory and domain-associated interactions preferentially contributed to the wild-type genome organization, whereas long-range interactions between repressive regions gained structural impact following cohesin loss (Fig. 3o). Together, these results show that cohesin depletion does not simply erase chromatin interactions but redistributes their organizational importance from local TADs and regulatory-element networks toward long-range compartmental and Polycomb-associated contacts.

### Structure-defining chromatin interactions are preserved across experimental modalities and genomic scales

We next asked whether the redistribution of structural importance observed in Hi-C data could also be recovered from an experimental modality that measures chromatin organization through a fundamentally different lens. We therefore applied HiCONA to Optical Reconstruction of Chromatin Architecture (ORCA)^58^ maps of the Sox2 locus in untreated and RAD21-depleted mESCs (Fig. 4a)^59^. Whereas Hi-C measures population-averaged ligation frequencies, ORCA reconstructs spatial distances between sequentially imaged genomic regions in individual cells^58^. The two approaches consequently produce distinct orthogonal information on 3D genome organization, providing an opportunity to test whether EIS extracts a shared organizational signature despite modality-specific measurements. In untreated cells, ORCA-derived EIS maps delineated the Sox2 TAD as a coherent region of structurally impactful interactions (Fig. 4b). Following RAD21 depletion, this organization became more fragmented, and contacts with high structural impact increasingly extended beyond the boundaries of the untreated domain (Fig. 4c). Differential analysis of raw ORCA contacts identified several interactions enriched after depletion that were not detectable in the corresponding Hi-C analysis, consistent with the distinct definitions of contact used by imaging and ligation-based assays. Despite these differences, differential EIS recovered the same directional transition in both modalities: interactions contributing preferentially to the untreated organization were concentrated within the Sox2 TAD, whereas those gaining importance after cohesin depletion extended across its former boundaries (Fig. 4d). Thus, the analyses indicated that Hi-C and ORCA can converge on the same structural interpretation even when their raw differential contact maps differed.

The ORCA-derived reorganization also involved contacts among Sox2 CREs. Enhancer–enhancer interactions exhibited significantly higher EIS values in untreated cells than in RAD21-depleted cells, consistent with the reduction observed in Hi-C data (Fig. 4e). Among high-impact contacts, both promoter–enhancer and enhancer–enhancer interactions were preferentially associated with the untreated condition. Annotation of differential EIS contacts by chromatin states further revealed distinct epigenetic compositions between the two conditions, indicating that the cohesin-depleted state was associated with differences in both the spatial distribution and regulatory identity of structurally impactful interactions. (Fig. 4f). These results show that the reduced contribution of enhancer-associated contacts and the emergence of boundary-spanning organization upon cohesion depletion represent emerging structural features depicted by HiCONA, irrespective of the adopted experimental modalities used to solve 3D genome organization.

We next investigated whether EIS could resolve structural reorganization at the finer spatial scale captured by Region Capture Micro-C (named RCMC)^30^ (Fig. 4g). RCMC detects localized, nested interactions named microcompartments, which frequently connect promoters, enhancers and CTCF-bound regions. We applied HiCONA at 1-kb resolution to an approximately 1-Mb region of chromosome 5 in untreated cells, following transcription inhibition or after RAD21 depletion^30^ (Fig. 4h–j). In untreated cells, annotated microcompartment contacts exhibited significantly higher EIS than the remaining interactions in the region, indicating that they contributed disproportionately to the inferred microscale organization (Fig. 4h, k). This enrichment persisted following transcription inhibition, consistent with microcompartments retaining structural importance in the absence of ongoing transcription, at least in this context^30^ (Fig. 4i, k).

Microcompartment contacts also retained significantly higher EIS than background interactions following RAD21 depletion, showing that they remained structurally distinguishable when loop extrusion was impaired (Fig. 4j, k). However, their EIS decreased relative to untreated cells, indicating that microcompartments persisted while becoming less influential within the alternative organization adopted by the depleted chromatin network. A similar pattern was recovered at an independent RCMC region on chromosome 8 (Extended Data Fig. 5a–e).

**Fig. 5.**
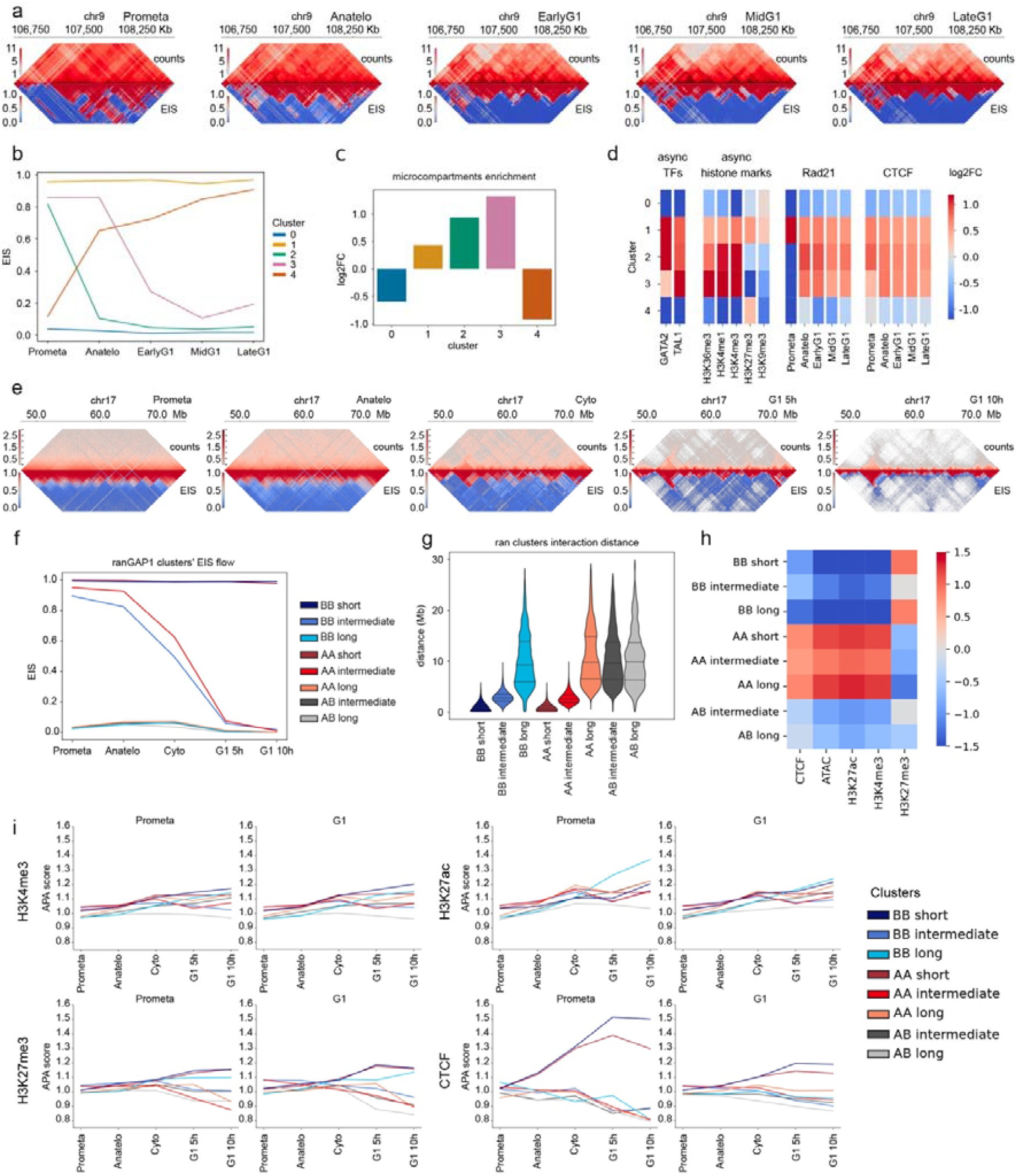
Entropy Impact Score reveals distinct trajectories of chromatin reorganization during mitotic exit a. Chromatin contact maps (top, log2 counts) and corresponding Entropy Impact Score (EIS) maps (bottom) for the genomic region chr9:106672000-108601600 at 1-kb resolution in G1E-ER4 erythroblasts. Maps are shown for prometaphase, anaphase–telophase, early G1, intermediate G1 and late G1. **b**. Median EIS trajectories throughout cell division for the 5 clusters identified by k-means clustering of contacts in G1E-ER4 erythroblasts. Number of contacts in each cluster: Cluster 0 = 352911, Cluster1 = 122581, Cluster2 = 49348, Cluster3 = 31080, Cluster4 = 36484. **c**. Log2 fold-change enrichment of microcompartment interactions across the five contact clusters in G1E-ER4 erythroblasts. Fold change is computed as fraction of microcompartments in the clusters divided by the fraction of microcompartment in the whole chromatin region of interest. **d**. Log2 fold-change enrichment of asynchronously bound transcription factors (GATA2 and TAL1), histone modifications (H3K36me3, H3K4me1, H3K4me3, H3K27me3, and H3K9me3), RAD21, and CTCF across the cell cycle in clusters identified using all chromatin contacts. **e**. Chromatin contact maps (top, log2 counts) and corresponding Entropy Impact Score (EIS) maps (bottom) for the genomic region chr17:47.5–73.4 Mb at 25-kb resolution in DLD-1 cells. Maps are shown for prometaphase, anaphase–telophase, cytokinesis, and G1 cells collected 5 h and 10 h after mitotic exit. **f**. Median EIS trajectories throughout cell division for the eight clusters identified by k-means clustering in untreated cells. Number of contacts in each cluster: BB_short = 575422, BB_intermediate = 254719, BB_long = 585956, AA_short = 492703, AA_intermediate = 368427, AA_long = 718256, AB_intermediate = 413735, AB_long = 650595. **g**. Distribution of chromatin interaction distances within DLD-1 untreated clusters. **h**. Log2 fold-change enrichment of CTCF, ATAC, H3K27ac, H3K4me1, and H3K27me3 CUT&RUN peaks across the untreated clusters. **i**. APA-score trajectories for H3K4me3 (top left), H3K27ac (top right), H3K27me3 (bottom left), and CTCF (bottom right) peaks in prometa and G1 phases in WT DLD-1 cells.

Together, these analyses show that structure-defining chromatin interactions can be resolved across experimental modalities and genomic scales. At the mesoscale, Hi-C and ORCA both revealed a redistribution of structural importance from local, enhancer-associated interactions toward distal contacts extending beyond the original domain boundaries. At the microscale, RCMC revealed that localized microcompartment interactions remained structurally distinguishable after cohesin loss, although their contribution to the surrounding organization was reduced. EIS therefore provides a common representation through which chromatin reorganization can be compared across ligation-based sequencing, spatial imaging and high-resolution Micro-C measurements, without requiring raw contact values.

### Structural trajectories define hierarchical programs of post-mitotic genome reassembly

Having established that EIS captures the redistribution of structural importance following defined perturbations, we next asked whether it could resolve the temporal reorganization of chromatin interactions during mitotic exit. We analyzed a Region Capture Micro-C time course spanning prometaphase, anaphase–telophase, early G1, mid G1 and late G1 across four genomic regions^31^ (Fig. 5a, Extended Data Fig. 6a-o). Rather than comparing chromosome-conformation maps independently at each stage, we followed variations in the EIS of individual chromatin interactions across successive cell-cycle states. We define the resulting evolution of their contribution to genome organization as a structural trajectory. K-means clustering of these trajectories identified five classes of contacts with distinct temporal dynamics, including interactions that remained structurally impactful throughout mitotic exit, interactions that rapidly lost importance after prometaphase and interactions that progressively gained importance during G1 (Fig. 5b). Microcompartment contacts were unevenly distributed across the five structural trajectories. Notably, cluster 1 combined microcompartment enrichment with persistently high structural importance throughout mitotic exit. In contrast, microcompartment-associated clusters 2 and 3 showed high EIS in prometaphase followed by a marked decline, whereas cluster 4 progressively gained structural importance during G1 and was depleted of microcompartment interactions (Fig. 5c). To determine whether distinct structural trajectories were associated with specific chromatin environments, we integrated ChIP–seq profiles for histone modifications with cell-cycle-resolved CTCF and RAD21 occupancy. Cluster 1 showed enrichment for both H3K4me3 and H3K27me3, consistent with a bivalent chromatin signature. Clusters 2 and 3 were enriched for active chromatin marks, including H3K36me3, H3K4me1 and H3K4me3, and showed preferential association with CTCF and RAD21 (Fig. 5d). By contrast, cluster 4 was comparatively depleted of active features and showed enrichment for the repressive mark H3K27me3. These results suggest that active, CTCF- and cohesin-associated contacts and microcompartment interactions carry substantial structural importance already in prometaphase but follow distinct trajectories during mitotic exit, whereas contacts that progressively acquire structural importance during G1 arise within a more repressive chromatin environment at a micro scale resolution.

**Fig. 6.**
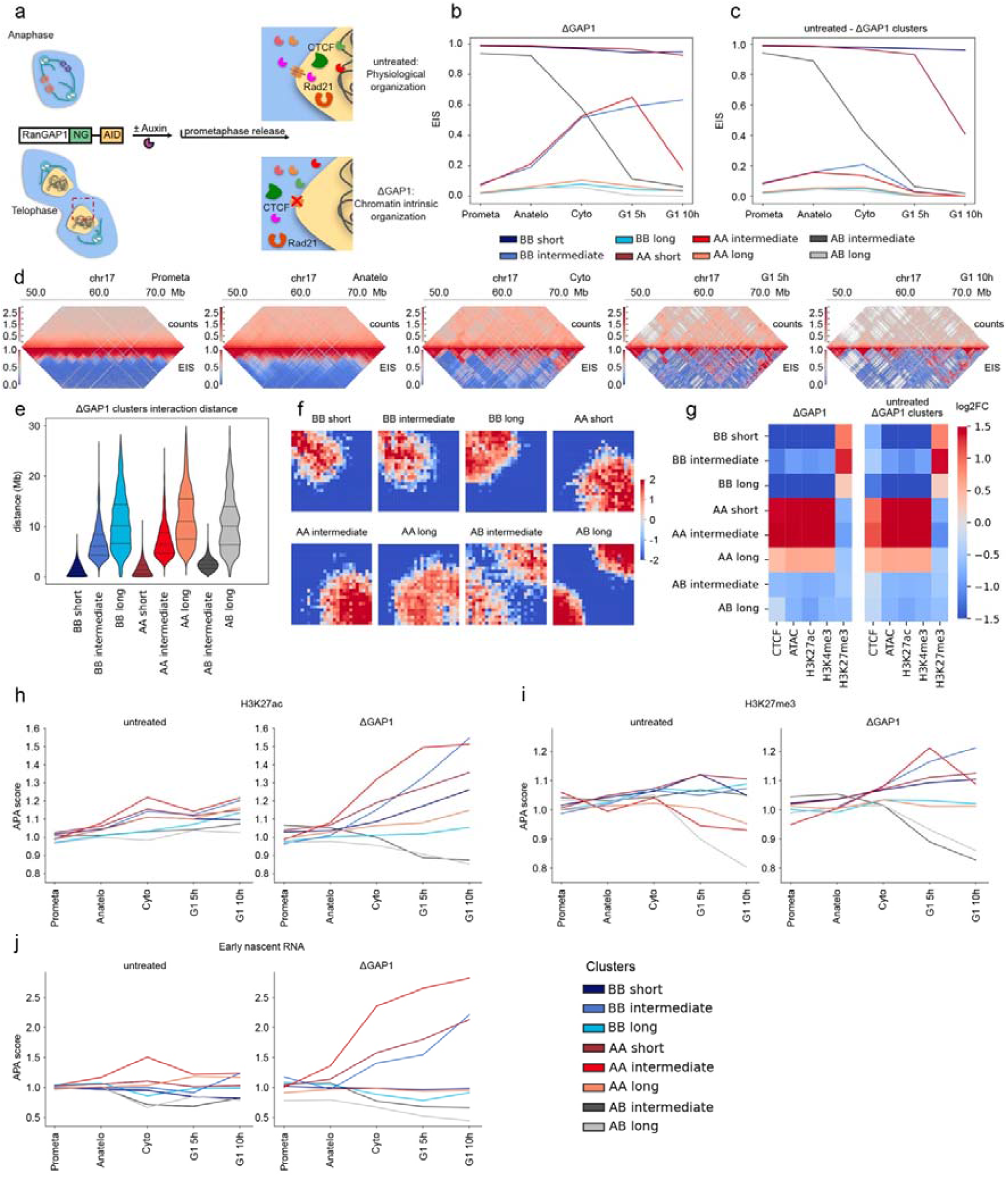
RNA-associated chromatin interactions define an intrinsic program of genome reorganization during mitotic exit a. Schematic representation of the effects of RanGAP1 deletion on protein nuclear import during the anaphase-to-telophase transition. **b**. Median EIS trajectories in ΔGAP1 condition throughout cell division for the eight clusters identified by k-means clustering in ΔGAP1 cells. Number of contacts in each cluster: BB_short = 355578, BB_intermediate = 213145, BB_long = 729470, AA_short = 929824, AA_intermediate = 859038, AA_long = 238732, AB_intermediate = 338448, AB_long = 395585. **c**. Median EIS trajectories in untreated condition throughout cell division for the eight clusters identified by k-means clustering in ΔGAP1 cells. **d**. Chromatin contact maps (top, log2 counts) and corresponding Entropy Impact Score (EIS) maps (bottom) for the genomic region chr17:47.5–73.4 Mb at 25-kb resolution in DLD-1 ΔGAP1 cells. Maps are shown for prometaphase, anaphase–telophase, cytokinesis, and G1 cells collected 5 h and 10 h after mitotic exit. **e**. Distribution of chromatin interaction distances within ΔGAP1 clusters. **f**. Saddle plots of subcompartment interaction enrichment for the clusters identified in ΔGAP1 cells. **g**. Log2 fold-change enrichment of CTCF, ATAC, H3K27ac, H3K4me1, and H3K27me3 CUT&RUN peaks across the ΔGAP1-derived clusters. Enrichment is shown for ΔGAP1 (left) and untreated (right) conditions. **h**. APA scores of chromatin contacts associated with H3K27ac peaks throughout cell division. APA scores are shown for untreated (left) and ΔGAP1 (right) cells, stratified by the clusters identified in ΔGAP1 cells. **i**. APA scores of chromatin contacts associated with H3K27me3 peaks throughout cell division. APA scores are shown for untreated (left) and ΔGAP1 (right) cells, stratified by the clusters identified in ΔGAP1 cells. *j*. APA scores of chromatin contacts associated with early nascent RNA genes throughout cell division. APA scores are shown for untreated (left) and ΔGAP1 (right) cells, stratified by the clusters identified in ΔGAP1 cells.

To resolve microcompartment-specific patterns of structural reorganization, we performed the temporal clustering using only annotated microcompartment contacts. This analysis identified five structural trajectories that largely recapitulated those observed across the complete interaction set (Extended Data Fig. 6p). The clusters characterized by persistently high EIS or by a progressive gain in EIS during G1 were enriched for shorter-range interactions, whereas microcompartments that lost structural importance after prometaphase generally spanned greater genomic distances (Extended Data Fig. 6q). Integration with chromatin annotations further showed distinct molecular identities among these structural trajectories. Notably, cluster 4 whose EIS increased during G1, showed a pronounced enrichment for H3K27me3 combined with a distinct increment of RAD21 and CTCF occupancy throughout cell cycle states (Extended Data Fig. 6r). Together, these analyses reveal that microcompartment reassembly is not uniform during mitotic exit. Instead, microcompartment contacts follow distinct structural trajectories associated with their genomic range and chromatin environment: short-range contacts preferentially retain or acquire structural importance during G1, whereas active and repressive microcompartments display separable patterns of post-mitotic reorganization.

To determine whether the structural trajectories identified at microcompartment scale extend to mesoscale chromosome organization, we analyzed a Hi-C time course dataset spanning prometaphase, anaphase–telophase, cytokinesis, early (5h), and late G1 (10⍰h)^33^. We applied HiCONA to nine different genomic regions spanning 20–30⍰Mb at 25-kb resolution (Fig. 5e and Extended Data Fig. 7). EIS maps revealed a transient expansion of structurally impactful interactions over broader genomic distances from prometaphase through cytokinesis, followed by an abrupt contraction of long-range structural interactions upon G1 entry (Extended Data Fig. 8). To identify the interactions underlying this transition, we classified contacts according to their structural trajectories, compartment association, and genomic distances. This approach resolved eight interaction classes comprising homotypic A/A or B/B, and heterotypic A/B contacts with distinct distance-dependent behaviors (Fig. 5f, g and Extended Data Fig. 9a-j). Short-range homotypic contacts within both A and B compartments retained high EIS throughout the time course. By contrast, intermediate-range homotypic interactions remained structurally important through cytokinesis but rapidly lost importance in G1, whereas long-range and heterotypic interactions generally maintained low EIS across the sampled stages. Thus, as observed at microcompartment scale, persistent structural importance during mitotic exit was preferentially associated with short-range interactions, while intermediate contacts contributed more transiently to chromosome organization.

To define the chromatin states of the mesoscale interaction classes, we integrated ATAC–seq and CUT&RUN profiles for histone modifications with CTCF occupancy. Contacts assigned to A compartments were enriched for accessible chromatin, CTCF binding, and active histone marks, whereas B compartment contacts were preferentially associated with H3K27me3 (Fig. 5h). Heterotypic A/B interactions showed no enrichment for any of the examined features (Fig. 5h). Saddle plots independently confirmed the expected compartment identities of the interacting anchors within each class (Extended Data Fig. 10e). We next asked whether the distinct structural trajectories could be accounted for their preferential interactions between loci sharing specific chromatin features. We reasoned that, if a given chromatin feature were a major determinant of a structural trajectory, contacts associated with that feature should display a corresponding stage-dependent change in interaction enrichment. To test this possibility, we performed aggregate peak analysis (APA) using H3K4me3-, H3K27ac-, H3K27me3- and CTCF-associated regions defined from chromatin profiles retrieved from cells in prometaphase or G1 and followed their interaction enrichment across the cell-cycle sampled stages (Fig. 5i and Extended Data Fig. 10a–d). Overall, contacts defined by the prometaphase and G1 chromatin states displayed broadly similar interaction profiles, with the most evident stage-related differences observed for CTCF-associated interactions defined in prometaphase. However, these differences did not recapitulate the distinct temporal patterns of structural importance identified by EIS. Thus, although chromatin accessibility and histone modifications distinguish the molecular identities of the inferred contact classes, the interaction dynamics associated with any single feature were insufficient to account for their distinct structural trajectories.

Together, these analyses identify a recurring distance-dependent principle of post-mitotic genome reassembly across micro- and mesoscale chromatin maps. Short-range homotypic contacts preferentially retained or progressively regained structural importance, whereas broader interactions contributed more transiently and declined rapidly after G1 entry. These trajectories occurred within distinct active and repressive chromatin environments but were not readily explained by contact frequency among any single assayed histone modification or architectural factor. This raised whether post-mitotic structural trajectories are predominantly established post-mitosis by trans factors such as chromatin regulators and RNA molecules or are partly guided by molecular and topological information retained within chromatin. We therefore analyzed the same cell cycle stages in cells in which nuclear import was impaired to identify structural trajectories that persist when the post-mitotic availability of trans-acting chromatin regulators is impaired^33^.

### Chromatin-intrinsic structural trajectories encode post-mitotic genome memory

To determine how trans-factors contribute to the structural trajectories identified during chromosome reassembly upon mitotic exit, we analyzed a DLD-1 model in which RanGAP1 depletion (ΔGAP1) impairs nuclear import upon re-establishment of the nuclear envelope during the anaphase-to-telophase transition^33^ (Fig. 6a). This perturbation reduces the post-mitotic availability of multiple *trans*-acting chromatin regulators, thereby allowing us to examine the contribution of *cis*-acting signals embedded within chromatin that persists when nuclear import is compromised^33^. We applied HiCONA to the same cell-cycle stages and genomic regions analyzed in untreated cells and classified interactions according to their EIS trajectories, genomic distances and compartment association. To specifically resolve the structural programs emerging when nuclear import is impaired, trajectory classes were defined using the ΔGAP1 time course (Fig. 6b and Extended Data Figs. 9 a-j). We then followed the same ΔGAP1-defined interaction classes in untreated cells, allowing direct comparison of their structural trajectories under perturbed and physiological conditions (Fig. 6c). Of note, because the interaction classes were defined from ΔGAP1 cells, the untreated trajectories in Fig. 6c differ from those in Fig. 5i, where clustering was independently defined from the time course of RanGAP proficient cells. This matched comparison revealed pronounced condition-dependent differences. Notably, ΔGAP1-defined intermediate-range homotypic A/A and B/B interactions progressively gained structural importance during mitotic exit and remained highly impactful into G1, whereas the same contacts showed only a transient increase in RanGAP1proficient cells before rapidly losing structural importance following cytokinesis (Fig. 6b, c).

Inspection of chromosome-wide EIS landscapes revealed the spatial organization underlying these altered trajectories. ΔGAP1 cells retained broad, compartment-like regions of high structural impact after cytokinesis and into G1, whereas this organization was progressively restricted upon G1 entry in RanGAP proficient cells (Fig. 6d and Extended Data Figs. 11, 12). Consistent with this chromosome-wide pattern, the ΔGAP1-defined intermediate A/A and B/B classes encompassed broader genomic distances than the intermediate classes independently defined in RanGAP proficient cells (Fig. 6e and Fig 5j). Thus, impaired nuclear import not only altered the temporal evolution of structural importance but also changed the genomic range of the interactions contributing to these trajectories. Saddle analysis further resolved the compartment-like organization of the ΔGAP1-defined classes, revealing distinct segregation patterns among homotypic A/A, B/B and heterotypic A/B interactions (Fig. 6f). We next asked whether this these pronounced structural differences reflected a corresponding change in the chromatin identity of the interacting regions. We therefore compared the enrichment of chromatin accessibility, CTCF and histone modifications within the ΔGAP1-defined classes between ΔGAP1 and proficinet cells. Despite their markedly different structural trajectories, the same interaction classes retained broadly comparable chromatin-state signatures in the two conditions: A/A classes remained enriched for active chromatin features, B/B classes for repressive H3K27me3-associated chromatin, and heterotypic classes showed no enrichment for either state (Fig. 6g). Thus, impaired nuclear import substantially alters the structural contribution and spatial extent of chromatin interactions without erasing the underlying epigenomic state of the interacting regions. Together, these results indicate that the post-mitotic remodeling of intermediate-range compartment-associated interactions is influenced by the nuclear import of trans factors rather than representing a consequence of chromosome decondensation upon mitotic exit. Indeed, in the absence of efficient nuclear import, a distinct subset of compartment-associated interactions remains structurally influential over broader genomic distances and persists into G1 (Extended Data Fig. 12).

Because ΔGAP1 cells retained stronger compartment-like organization despite the broadly comparable chromatin-state identity of the corresponding interaction classes, we asked which molecular features already present on mitotic chromosomes characterize the genomic loci underlying these altered structural trajectories. We first examined whether chromatin state was associated with changes in interaction strength during mitotic exit, using APA to track contacts frequency between genomic regions carrying selected histone modifications along the sampled stages. Homotypic H3K27ac-associated contacts showed a progressive increase in interaction strength in ΔGAP1 cells, emerging predominantly from cytokinesis and remaining elevated through G1, whereas the corresponding increase in proficient cells was substantially reversed (Fig. 6h, and Extended Data Fig. 13a). H3K27me3-associated contacts displayed a distinct temporal behavior. Their major changes emerged predominantly after cytokinesis, and differed across structural trajectories, with selected interaction classes gaining interaction strength during G1 while others progressively declined (Fig. 6i, and Extended Data Fig. 13b). Thus, active and repressive chromatin states were associated with distinct patterns: H3K27ac-associated interactions showed an early and progressive reinforcement following anaphase–telophase that was prolonged by impaired nuclear import, whereas changes involving H3K27me3-associated regions emerged predominantly after cytokinesis and were more trajectory-specific.

Given the early and progressive behavior of H3K27ac-associated contacts, we next asked whether this pattern was related to genomic regions that remain transcriptionally competent during mitosis. We therefore used nascent transcription measured in prometaphase to define mitotically transcribed regions and followed their interaction strength throughout mitotic exit. Strikingly, contacts involving prometaphase-transcribed regions displayed a pattern closely resembling that observed for H3K27ac-associated chromatin: their interaction strength increased in ΔGAP1 cells from anaphase–telophase and continued to rise through cytokinesis into G1, particularly within the A/A intermediate, and A/A short classes (Fig. 6j and Extended Data Fig. 13c). In untreated cells, the same interaction sets initially increased through cytokinesis but subsequently declined upon G1 entry. Thus, genomic regions transcriptionally active in prometaphase participate in a transient increase in interaction strength during mitotic exit, whereas this configuration is prolonged into G1 when post-mitotic nuclear import is impaired.

The similarity between the interaction dynamics of H3K27ac-associated and mitotically transcribed regions raised the possibility that these two chromatin-intrinsic features jointly characterize the contacts showing the strongest post-mitotic reinforcement. We therefore stratified prometaphase-transcribed loci according to their histone-modification status. The strongest increase in interaction strength occurred between regions that were both transcriptionally active and H3K27ac-positive in prometaphase (Extended Data Fig. 13d). This increase exceeded that observed for prometaphase-transcribed regions associated with H3K4me3 or for contacts defined by H3K27ac status in G1 (Extended Data Fig. 13e, f). Conversely, the progressive increase in interaction strength was lost when prometaphase-transcribed regions lacking H3K27ac were considered (Extended Data Fig. 13g). Thus, the combination of mitotic transcriptional activity and H3K27 acetylation most strongly characterize genomic regions whose interactions remain reinforced following mitotic exit when nuclear import is impaired.

We finally asked whether the genes contributing to this mitotic transcriptional program reflected a DLD-1-specific state. Gene ontology analysis revealed highly similar functional categories among mitotic nascent transcripts detected in proficient and ΔGAP1 DLD-1 cells, which were also shared with the liver cancer Huh7 cells^60^ (Extended Data Fig. 13h, i and Extended Data Tables 1-3). Notably, both the shared and the cell type-enriched GO terms refers to gene categories related to homeostatic cellular functions such as regulation of gene expression, chromatin factors, and mitosis regulation.

Together, these results identify molecular features benchmarking mitotic chromosomes that characterize genomic regions contributing to the altered structural trajectories revealed upon impaired nuclear import. Active and repressive chromatin states display distinct interaction dynamics, with the progressive behavior of H3K27ac-associated contacts closely paralleling that of mitotically transcribed regions. Their intersection identifies a subset of transcriptionally competent, H3K27ac-marked mitotic chromatin whose interaction strength increases during mitotic exit and remains elevated into G1 when nuclear import is impaired. These regions therefore represent a candidate chromatin-intrinsic substrate associated with the structural information retained during post-mitotic genome reassembly.

## Discussion

In this work we address a fundamental conceptual paradox related to mitosis representing an obstacle for faithfully propagating the epigenetic memory while every cell division reconstructs both chromatin states and three-dimensional genome organization, ensuring the activation of the cell-type specific transcriptional program ^2 18 19^. We therefore investigated whether mitosis itself could contribute to cellular memory by preserving structural information that guides genome refolding in daughter cells. We found that structure-defining chromatin interactions that evolve along cell cycle states delineate chromatin-intrinsic structural trajectories, providing a candidate structural basis for post-mitotic genome memory. Our data link mitosis-specific configurations to subsequent folding, with active and repressive chromatin states contributing through distinct temporal trajectories. Notably, prometaphase-transcribed regions marked by H3K27ac emerge as chromatin-intrinsic domains retaining molecular and topological information that may guide genome reassembly in daughter cells.

HiCONA provides a quantitative framework for investigating chromatin organization across multiple architectural scales. Conventional analyses characterize domains, compartments, and regulatory interactions using distinct metrics^43 44 45,50^. HiCONA combines an inferred hierarchical representation of genome organization with the Entropy Impact Score (EIS), which quantifies each contact’s relative contribution to that organization. This integration identifies hierarchical interaction patterns without predefined domain annotations and evaluates contacts associated with different architectural features within a common quantitative framework^44 45,50^. The advance lies in coupling multiscale structural inference to contact-level importance, enabling individual interactions to be followed as their contributions to genome organization change across perturbations or cellular states.

We benchmarked the value of this approach by determining the effects of acute depletion of Cohesin on the changes in contacts structural importance across genomic scales. RAD21 loss reduced the structural contribution of TAD-associated interactions and redistributed contact importance, reshaping local regulatory organization while increasing the contribution of long-range compartment-associated contacts. Long-range interactions between H3K27me3-enriched regions gained structural importance, highlighting Polycomb-associated contacts in the organization that emerges after cohesin loss^55^. HiCONA thus extends the established observation that cohesin depletion weakens TADs and enhances compartmentalization^37 21^ by resolving which contacts increasingly contribute to the reorganized genome folding^37 21^.

This redistribution may help explain why extensive architectural disruption can coexist with comparatively modest transcriptional changes. At the Sox2 locus, loss of loop extrusion perturbs TAD organization and distal Sox2–SCR contacts, while transcription remains resilient through regulatory mechanisms involving promoter-proximal elements and the distal enhancer cluster^56 57^. Our findings suggest that such functional robustness can coexist with extensive architectural reorganization because loss of TAD-scale structural importance is accompanied by a redistribution toward interactions operating at different genomic scales. Local regulatory interactions may therefore preserve the connectivity required for transcriptional output, while alternative longer-range interactions increasingly contribute to the organization of the surrounding chromatin landscape. In this framework, contact frequency, structural importance and regulatory function represent related but non-equivalent properties of chromatin interactions, and their separation may provide a means to understand how transcriptional programs remain buffered despite profound changes in higher-order genome architecture.

Beyond this functional interpretation, our analyses also revealed that the redistribution of structural importance varied with the genomic context over which chromosome organization was interrogated. Analysis of narrow genomic windows preferentially emphasized changes in local domain and regulatory organization, whereas extending the analysis to mesoscale revealed an additional layer of long-range organization. These observations suggest that the structural importance of a chromatin contact may not be determined by the interaction alone, but by its relationship with the broader organizational landscape in which it is embedded. This context-dependent view provides a potential framework for understanding how local and long-range chromatin interactions are reorganized following perturbation. Future studies combining HiCONA with orthogonal structural and functional approaches will be required to establish the biological relevance of these context-dependent relationships and determine how interactions identified as structurally important at different genomic scales contribute to chromosome organization and function.

Studies of post-mitotic genome reassembly have described distinct kinetics for the recovery of local contacts, cohesin-dependent domains and compartmentalization. HiCONA adds a complementary perspective by following the structural contribution of individual interactions across these transitions. Across micro- and mesoscale organization, short-range homotypic interactions preferentially retained or progressively regained structural importance, whereas longer-range interactions contributed transiently before declining in importance upon G1 entry. These trajectories were associated with distinct chromatin environments, encompassing active, repressive and microcompartment-associated contacts and were already distinguishable during mitosis. Compartment-associated interactions could carry substantial structural importance before compartmentalization became evident, while microcompartment-associated contacts followed divergent trajectories. Thus, the time at which an architectural feature becomes detectable does not coincide with the onset of structural contributions from its constituent interactions. The recovery of interphase architecture may therefore reflect the emergence of recognizable structures from partially overlapping trajectories, rather than a strictly ordered progression through genomic scales.

Overall, our findings extend the concept of mitotic bookmarking from the preservation of local molecular states to the emergence of chromatin-intrinsic structural trajectories, suggesting that epigenetic information retained on mitotic chromosomes may bias the path through which three-dimensional genome organization is reconstructed in daughter cells. We propose that post-mitotic genome organization follows a hysteretic rather than a hierarchical folding process. In this framework, chromatin architecture depends on the structural trajectory established before and during mitosis. Rather than representing a transient loss of organization, the mitotic chromosome constitutes an informative intermediate state that biases subsequent folding. Our data suggest that transcriptionally competent euchromatic regions, maintained through mitotic bookmarking and engaging in homotypic interactions function as architectural nucleation sites that preserve a chromatin-intrinsic folding trajectory. In the absence of nuclear import, these intrinsic trajectories continue to strengthen, indicating that transfactors primarily tune post-mitotic folding. Structural trajectories guiding genome reassembly provide structural inertia that buffers transient perturbations and preserves epigenetic memory, whereas sustained regulatory inputs and nuclearimport–dependent factors can redirect the trajectory, enabling plasticity during the mitotic exit window.

## Methods

### Processing of Hi-C, Micro-C and RCMC data

Publicly available Hi-C, Micro-C and Region Capture Micro-C (RCMC) contact matrices were analyzed in the form in which they were deposited by the original studies. Because the analysis started from contact matrices rather than sequencing reads, no read alignment, filtering, deduplication, or contact-matrix construction was performed. For each dataset, the relevant genomic regions and matrix resolutions were selected as reported in Supplementary Table 5. HiCONA was applied to the unbalanced contact counts stored in the contact matrices. No matrix balancing or observed-over-expected normalization was applied before network construction.

### Compartment analysis and peak calling

Subcompartment classification of GM12878, IMR90, and DLD-1 (WT and ΔGAP1) cells we relied on Calder2^61^. The resulting subcompartment annotations were mapped to the genomic bins used in the corresponding downstream analyses.

Peak calling for chromatin-profiling datasets was performed using workflows implemented in Nextflow^62^. For ATAC-seq and CUT&RUN data, the nf-core pipeline atacseq (v2.1.2)^63^ was run with default parameters on hg38 genome (GRCh38.114). For datasets without exogenous spike-in controls, peaks were called using MACS2^64^. When spike-in controls were available, spike-in normalization and peak calling were instead performed using SpikeFlow^64^. The resulting peak coordinates were used for the genomic annotation and enrichment analyses described below.

### Processing of ORCA imaging data

We processed the ORCA datasets following the procedure described in the original publication^59^. Each dataset consists of a set of matrices in which every DNA probe is annotated with its three-dimensional position in Cartesian space (x, y, z). Each matrix corresponds to an independent experimental replicate. For each replicate, we first computed the Euclidean distance between all probe pairs. To convert spatial distances into binary contact maps, we applied the thresholding criterion defined by Hafner *et al*.: probe pairs separated by less than 256 nm were classified as interacting. For each pair of probes, the total number of contacts was then obtained by counting the number of replicates in which the distance fell below this threshold^59^.

### Chromatin network construction

Chromatin networks were generated from chromatin contact maps, in which the genome is partitioned into genomic regions of equal length, known as bins. The number of contacts between two bins represents the total number of paired reads mapped to those genomic regions.

To initiate chromatin network construction, a specific region of interest must be selected. Each bin within this region is represented as a node in the network, and an edge is established between two nodes if at least one chromatin contact has been detected between the corresponding bins.

Let *V* = {*1,2*,…,*n*} be the set of nodes of the chromatin network and *E* = {*e*_*12*_, *e*_*1n*_,…*e*_*nm*_} the set of edges linking the nodes. Chromatin network weights *W* = {*w*_*12*_, *w*_*1n*_,…*w*_*nm*_*}* are modelled as follows:

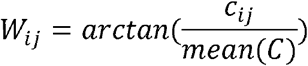

Where *c*_*ij*_ is the number of contacts of the edge *e*_*ij*_, connecting nodes *i* and *j*, and *C = {c*_*12*_, *c*_*1n*_,…*c*_*nm*_} is the set of non-zero edges’ counts.

### Nested stochastic block-model inference

The hierarchical organization of each chromatin network was inferred using a degree-corrected nested stochastic block model (nSBM), as implemented in graph-tool 2.95^51^. We supplied the arctangent-transformed contact weight for each observed edge to the model as a real-valued edge covariate through the recs argument and modeled it using the real-normal covariate type. No constraints based on linear genomic position or cluster contiguity were imposed during inference.

We inferred hierarchical partitions using minimize_nested_blockmodel_dl, which identifies the nSBM state that minimizes the total description length of the network and its hierarchical partition. This minimum-description-length criterion balances the ability of the model to represent the observed network against the additional complexity introduced by the inferred partition. The number of multilevel MCMC sweeps was set using multilevel_mcmc_args={“niter”: 10}.

Because the optimization procedure is stochastic, inference was repeated ten times from independent initializations. The state with the lowest description length was retained and subsequently refined by simulated annealing using mcmc_anneal. The inverse temperature beta was increased logarithmically from 1 to 10 over 1,000 annealing steps. At each value of beta, ten multiflip MCMC iterations were performed by setting force_niter=10, allowing the hierarchical partition to be refined through node-move and merge–split proposals. This stage was used exclusively to optimize the inferred hierarchy; no posterior samples were collected.

Hierarchical block assignments were extracted from the final NestedBlockState. Level 1 was defined as the finest inferred partition, and successively higher levels represented progressively coarser groupings of genomic bins. The terminal level containing a single block was excluded from downstream analyses.

### Entropy Impact Score

The Entropy Impact Score (EIS) was computed using the uncertain-network reconstruction framework implemented in UncertainBlockState in graph-tool^51^. MeasuredBlockSate or MixedMeasuredBlockSate are the preferred models when the maximum number of contacts that can occur between two bins is known, as is the case with ORCA data, where this corresponds to the total number of cells analyzed. For each chromatin network, the contact count *c*_*i,j*_ associated with an observed edge was transformed into an input edge probability *q*_*ij*_, scaling the arctan-transformed counts between 0 and 1 as follows:

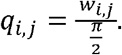

These values represent contact-derived estimates of the probability that the corresponding edges are present in the latent chromatin network.

For a simple undirected network containing N nodes and m observed edges, all node pairs not connected in the input network were assigned the same probability

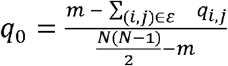

where ε denotes the set of observed edges. This definition ensures that the expected number of edges under the input probability model equals the number of edges in the observed network:

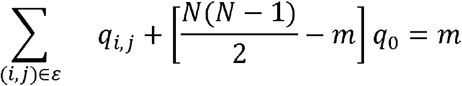

The edge-specific probabilities *q*_*ij*_ and the non-edge probability *q*_O_ were supplied to UncertainBlockState as q and q_default, respectively. A nested stochastic block model was used as a structural prior. The procedure jointly sampled the latent adjacency matrix *A* and its hierarchical block partition *b* from the posterior distribution *P(A,b* | *Q*), thereby integrating the contact-derived edge probabilities with the large-scale organization inferred from the network.

Before marginal collection, the Markov chain was equilibrated for 10,000 sweeps. Posterior edge marginals were then collected over 500,000 MCMC sweeps. The callback was applied after every 10 sweeps, yielding S=50,000 sampled latent networks.

For each node pair (i,j)(i,j), the EIS was defined as its posterior marginal probability of being present in the latent network:

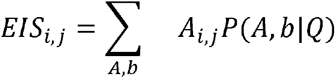

Accordingly, EIS values range from 0 to 1. Higher values indicate that an interaction is more consistently supported across the posterior ensemble after integrating its contact-derived input probability with the hierarchical structure of the network. For downstream analyses, EIS values were retained for all node pairs represented in the posterior marginal graph.

### Polymer simulations

To generate polymer models, we used the Polychrom Python library ^52^. We produced 20 in-silico Hi-C contact maps by varying the intensities of loop extrusion and compartmentalization. Each model was initialized with 4,000 monomers, representing a 10 Mb chromatin region. The mean size of each subdomain was 444 monomers (≈ 1.1 Mb), while the mean size of the 11 simulated TADs was 267 monomers (≈ 666.5 Kb).

To modulate homotypic compartment interactions, we simultaneously varied the HS parameters for the A and B compartments from 0 to 0.03 (in increments of 0.01) and from 0 to 0.3 (in increments of 0.1), respectively. To modulate loop extrusion strength, we varied the release probability (RP), using values of 0.0005, 0.005, 0.05, 0.5, and 1, while keeping the capture probability fixed at 0.9 and the loop extruder separation and lifetime at 800 and 200, respectively. For the simulation of the 30-Mb chromatin region, the HS parameters for the A and B compartments were set to 0.2 and 0.3, respectively, the RP was set to 0.04, the capture probability to 0.85, and the loop extruder separation and lifetime to 130 and 50, respectively.

### Validation against simulated structures

To evaluate whether HiCONA recovered the structural features encoded in the polymer models, each simulated contact map was converted into a chromatin network and analyzed using the same network-construction and hierarchical-inference procedures applied to the experimental datasets. Validation was performed across all 20 combinations of homotypic interaction strength (HS) and loop-extruder release probability (RP).

Recovery of loop-extrusion domains was assessed using the Adjusted Rand Index (ARI). The positions of the simulated loop-extrusion boundaries were first converted into a ground-truth partition by assigning each polymer monomer to its corresponding extrusion domain. For each non-trivial level of the HiCONA hierarchy, the vector of inferred cluster assignments was compared with the ground-truth domain assignments using ‘adjusted_rand_score’ from scikit-learn. An ARI of 1 indicates identical partitions, whereas values close to 0 indicate agreement comparable to that expected by chance. Negative values indicate agreement lower than expected by chance. ARI values were calculated independently for every hierarchical level and for each combination of HS and RP parameters.

Recovery of the simulated A/B compartment organization was quantified using the Gini impurity. For each HiCONA cluster *c*, the proportions of monomers assigned to compartments A and B were denoted by *p*_*c,A*_ and *p*_*c,B*_, respectively. The Gini impurity of the cluster was calculated as

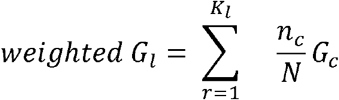

where *l* is the hierarchy level, r represents the index of a single cluster within a level, *K* _*l*_ is the total number of clusters in a single level, *n*_c_ is the number of monomers of a single cluster, and N is the total number of monomers in the polymer model.

A value of 0 indicates that all monomers within the cluster have the same compartment identity, whereas the maximum value of 0.5 indicates equal proportions of A and B monomers. Cluster-level Gini impurity values were summarized at each hierarchical level using the mean weighted by cluster size. This analysis was performed independently for every hierarchical level and for each combination of HS and RP parameters.

### TAD-calling benchmark

To evaluate HiCONA’s performance as a TAD caller, we used TADshop^53^, a platform that provides a curated set of TAD annotations obtained from 39 gold-standard TAD-calling algorithms. TADshop offers a preprocessed .cool file from GM12878 Hi-C data, which we used to ensure consistency with existing benchmarked methods.

The platform employs several metrics to assess TAD-calling performance, including: enrichment of structural proteins (e.g., CTCF, RAD21, SMC3) at TAD bondaries, differential enrichment of histone marks, specifically H3K36me3 vs. H3K27me3, insulation score at TAD borders, and contact enrichment within versus between TADs. For each method, TADshop normalizes individual metric values between 0 and 1, then computes an average score across metrics to derive a global performance ranking.

For benchmarking HiCONA, we used only the clusters identified at the first hierarchical level, corresponding to TAD-like structures.

### Genomic feature annotation

Promoters and enhancers were identified using ChIP-seq datasets for the histone modifications H3K27ac, H3K4me3, and H3K4me1. Active promoters were defined as transcription start sites (TSS) ± 500 bp regions that overlapped with both H3K27ac and H3K4me3 peaks. Enhancers were defined as genomic regions marked by the co-occurrence of H3K27ac and H3K4me1 peaks.

Genomic bins were annotated with binary labels based on the presence or absence of ChIP-seq peaks, promoter regions, or enhancer regions. A bin was assigned a score of 1 if it contained the feature of interest (e.g., promoter, enhancer, or specific histone mark), and a score of 0 otherwise.

For each replication timing stage (G1b, S1, S2, S3, S4, and G2), we aggregated read counts across genomic intervals corresponding to the Hi-C resolution used in the analysis, summing all reads mapping within each bin.

Compartment annotations were obtained from publicly available datasets and directly integrated into our analysis. Each chromatin bin was assigned a label corresponding to its compartment identity (A or B), based on the supplied annotations.

Microcompartment annotations were obtained from a manually curated .bedpe file provided by the laboratory that first described them^30^. This file contains genomic coordinates describing pairs of interacting anchors associated in a microcompartmental contact. To annotate chromatin contacts in our Hi-C matrices, we used the BEDTools suite^65^. A contact was classified as a microcompartment interaction if both bins forming the contact overlapped with the respective anchor regions from the .bedpe file.

### Replication-timing analysis

We analyzed replication-timing profiles independently across four levels of the HiCONA hierarchy. For each hierarchical level, we summarized Repli-seq signals across the genomic bins assigned to each cluster for six cell-cycle fractions: G1b, S1, S2, S3, S4, and G2. Let *B*_*c*_ denote the set of genomic bins assigned to cluster *c*, and let *n*_*c*_ = |*B* _*c*_| denote the number of bins in that cluster. The mean Repli-seq signal of cluster *c* in cell-cycle fraction *i* was calculated as

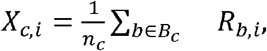

where *R*_*b,i*_ is the Repli-seq signal measured in genomic bin *b* during fraction *i*.

For each hierarchical level, the resulting matrix contained one row per HiCONA cluster and six columns corresponding to the six Repli-seq fractions. Principal component analysis was performed directly on this matrix using the PCA implementation in scikit-learn with two components. The variables were centered internally by the PCA algorithm, but no variance scaling was applied before decomposition. Each cluster was annotated according to the cell-cycle fraction exhibiting the highest mean Repli-seq signal:

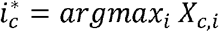

To quantify replication-timing enrichment while controlling for cluster size, the observed mean profile of each cluster was compared with profiles obtained from size-matched random genomic sets. For each cluster *c*, 1,000 sets containing *n*_c_ bins were sampled without replacement from the complete set of genomic bins included in the corresponding dataset. For randomization *r*, the mean Repli-seq signal was calculated separately for each fraction as

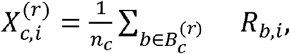

Where 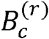 denotes the set of bins selected in randomization *r*. The mean and standard deviation of the 1,000 randomized profiles were then used to calculate a size-corrected Z-score:

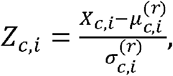

Where 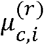 and 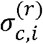 are the mean and standard deviation of the randomized mean signals for cluster *c* and fraction *i*, respectively.

The six Z-scores were combined into a Replication Timing Index (RTI) using weights representing the temporal order of the Repli-seq fractions:

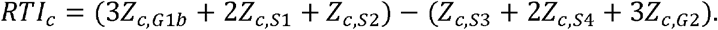

Higher RTI values therefore indicate preferential enrichment in early-replicating fractions, whereas lower values indicate enrichment in late-replicating fractions. At each hierarchical level, clusters with an RTI greater than the mean RTI across clusters were classified as early replicating, whereas clusters with an RTI equal to or below the mean were classified as late-replicating.

### Differential EIS analysis

Differential analyses were performed independently on Hi-C and ORCA datasets by comparing wild-type and cohesin-depleted contact maps covering the same genomic region at the same resolution. HiCONA was applied independently to each condition, and the EIS was calculated for every contact as described above. Corresponding bin pairs were aligned according to their genomic coordinates, and the differential EIS was calculated as

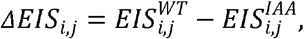

where *i* and *j* identify the two genomic bins forming the contact. Thus, positive values indicate contacts with higher EIS in WT cells, whereas negative values indicate contacts with higher EIS following cohesin depletion. Contacts with EIS values below −0.75 or above 0.75 were classified as differential EIS.

An analogous differential analysis was performed using contact frequencies. To account for differences in total contact coverage between conditions, the count of each contact was divided by the sum of the counts of all contacts retained within the corresponding genomic region:

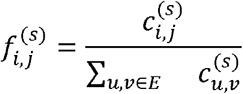

where 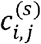 is the contact count between bins *i* and *j* in condition *s*, and the denominator is the total contact count within the analyzed region in that condition. The differential normalized contact frequency was then calculated as

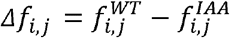

Positive values indicate contacts relatively enriched in WT cells, whereas negative values indicate contacts relatively enriched in cohesin-depleted cells.

### Temporal EIS trajectory clustering

To identify chromatin contacts exhibiting similar temporal changes in structural importance, HiCONA was applied independently to contact maps obtained at each cell-cycle stage, and the EIS was calculated for every contact as described above. For each dataset, contact maps covering the same genomic regions at the same resolution were used, and corresponding contacts were aligned across stages according to their genomic coordinates. For the G1E-ER4 RCMC dataset, each contact was represented by a five-dimensional vector containing its EIS values in prometaphase, anaphase–telophase, early G1, intermediate G1 and late G1. For each contact between bins *i* and *j*, a temporal EIS vector was constructed as

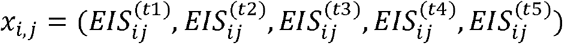

where *t*1, *t*2, *t*3, *t*4, *t*5 denote the ordered cell-cycle stages.

For the DLD-1 Hi-C dataset, each contact was instead represented by 15 features: five EIS values, five mean CALDER2 subcompartment scores and five standard deviations of the CALDER2 scores, with one value of each type for each cell-cycle stage. Let 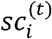 and 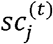 denote the continuous CALDER2 subcompartment scores assigned to the two anchors of contact *i, j* at stage *t*. Their mean was calculated as

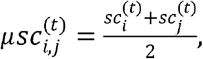

and their variability was summarized as

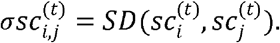

The complete DLD-1 feature vector was therefore

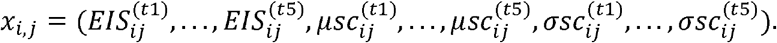

The five DLD-1 stages were ordered as prometaphase, anaphase–telophase, cytokinesis, G1 early and G1 late.

Temporal profiles were grouped using k-means clustering. For both datasets, the number of clusters was determined using the elbow method. K-means was fitted across a range of candidate *K* values, and the within-cluster sum of squares was calculated as

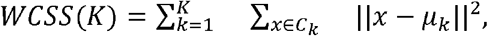

where *C*_*k*_ is the set of contacts assigned to cluster *k* and *µ*_*k*_ is its centroid. The value of *K* corresponding to the elbow of the WCSS curve, beyond which additional clusters produced only limited reductions in WCSS, was selected. This procedure identified five clusters for the G1E-ER4 dataset and eight clusters for the DLD-1 dataset.

For the DLD-1 comparison, clustering was performed using the profiles measured in *Δ*GAP1 cells. The resulting contact assignments were retained when summarizing the corresponding untreated contacts. The representative EIS trajectory of each cluster was calculated as the median EIS of its contacts at each stage:

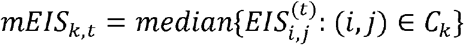

### Aggregate peak and saddle analyses

Aggregate peak analysis (APA) was performed using cooltools^66^. For each set of genomic interactions, cis-expected contact frequencies were calculated from the balanced contact matrix using cooltools.expected_cis. Observed-over-expected contact windows were subsequently extracted around the corresponding interaction coordinates using cooltools.pileup. A flank of 500 kb, corresponding to 20 bins on either side of the interaction at 25-kb resolution, was used to generate a (41×41)-pixel window for each contact. Individual windows were averaged element-wise using numpy.nanmean, thereby excluding missing or invalid pixels from the calculation.

Interaction enrichment was quantified by comparing the central pixel of the aggregate contact map with its local background. The background was defined as a donut-shaped region comprising pixels located at a Euclidean distance of 5–10 pixels from the center. The APA enrichment score was calculated as

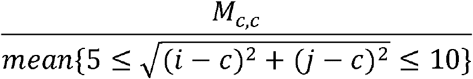

where (M) denotes the aggregate observed-over-expected contact matrix and (c) its central pixel. APA scores greater than 1 indicate that the analyzed interaction anchors exhibit greater contact enrichment than their surrounding local background.

Saddle plots were generated to characterize the subcompartmental context of cis interactions assigned to each EIS trajectory cluster. For each cell-cycle stage and experimental condition, genomic bins were annotated with continuous subcompartment scores derived using CALDER2^61^. Scores were oriented from the most B-like to the most A-like chromatin state and rank-transformed to values ranging from 0 to 1. Ranked genomic bins were then divided into 25 equal-width intervals. Balanced Hi-C contact frequencies were normalized by the expected contact frequency at the corresponding genomic separation, and the resulting observed-over-expected values were averaged within each pair of subcompartment-score intervals to generate a two-dimensional saddle matrix. Missing values were excluded from the calculation. Values above and below 1 indicate contact enrichment and depletion, respectively, relative to loci separated by comparable genomic distances. Saddle plots were generated separately for each cell-cycle stage and for the untreated and ΔGAP1 conditions using DLD-1 Hi-C matrices at 25-kb resolution. The analysis included all genomic regions for which both EIS trajectory assignments and subcompartment annotations were available.

### Running Time

HiCONA runtimes on datasets with different numbers of contacts and different coverages are present in Supplementary Table 6.

### Statistical analysis

To assess the performance of HiCONA in identifying chromatin topologies, we employed the following metrics and tests. Adjusted Rand Index (ARI) was used to evaluate the accuracy of TAD detection in polymer model simulations by comparing HiCONA’s hierarchical clusters with ground truth TAD assignments.

Gini Index was applied to quantify the purity of HiCONA-derived clusters in terms of compartment (A/B) identity, both in polymer models and experimental datasets.

One-sided Mann–Whitney U tests were used throughout the analysis to determine statistical significance when comparing EIS values, cluster metrics, or other distributions across different conditions (e.g., WT vs. cohesin depletion or transcription inhibition).

All statistical analyses were performed using standard libraries in Python, and p-values below 0.05 were considered statistically significant. For visualization purposes, significance levels were represented as follows:

⍰ p < 0.05: ^*^

⍰ p < 1 × 10^−3^: ^**^

⍰ p < 1 × 10^−5^: ^***^

## Data availability

All datasets used in this project were retrieved from publicly available sources.

GM12878 and IMR90 Hi-C data^23^ are available as processed files at multiple resolutions through the 4D Nucleome data portal (4DNFIXP4QG5B and 4DNES1ZEJNRU). For TAD benchmarking, we used the same cell type and experimental dataset of GM12878 cells, but processed according to the workflow described in^53^ and obtained via TADshop (TADshop.unil.ch).

For perturbation analyses, we focused on cohesin depletion and transcription inhibition in mESCs, integrating datasets from two different sources. High-resolution (1 kb) chromatin interaction data were obtained from Region Capture Micro-C (RCMC) experiments^30^ (GSE207225), while datasets centered on the *Sox2* locus were derived from complementary Hi-C experiments^55^ (E-MTAB-7816). Microcompartment interactions^30^ are available at https://github.com/ahansenlab/RCMC_analysis_code in .bedpe format.

ORCA data used for the comparison of HiCONA performance with HiC data at Sox2 locus were retrieved from 4D Nucleome Data Portal, both for wild-type (4DNESLRTOSQT) and cohesin-depleted (4DNESBH54BG2) conditions.

RCMC cell cycle dataset^31^ can be found at GEO database (GSE276657). While the Hi-C cell cycle dataset WT and ΔGAP1^33^ can be found at GEO database (GSE277875).

Repliseq scores were retrieved from ENCODE (ENCSR218XPB) for each cell cycle stage. ChIP-seq track for mESC were retrieved from ENCODE: H3K27ac (ENCSR000CGQ), H3K4me1 (ENCSR000ADK), and H3K4me3 (ENCSR000ADL). H3K27me3 Chip-seq tracks were retrieved from Gene Expression Ominbus (GSE98671). Data used to characterize cell cycle Hi-C data were obtained by the original manuscript^33^: ATAC-seq (GSE277731), CUT&RUN (GSE308844), and SLAM-seq (GSE309609).

## Code availability

The software of HiCONA can be found on github (https://github.com/leomorelli/HiCONA). While the script used for benchmarking and testing HiCONA on chromatin organization data can be found at (https://github.com/leomorelli/HiCONA_code).

## Acknowledgement

We would like to thank Marco Di Stefano (IGH, University of Montpellier) and Elisa Piperni (University of Trento) for the discussions and the precious hints. We also would like to thank Sarah D’Annunzio (Ludwig-Maximilians-Universität München) and Giacomo Costalunga (Max Planck Institute for Biological Intelligence) for their support in data visualization. Leonardo Morelli’s work was supported by AIRC Post-doctoral fellowships (#3140) and Short-EMBO Fellowship (#11909). Work in the Zippo group was supported by Italian Association for Cancer Research (AIRC) IG# 2291 and IG# 30745, and Worlwide Cancer Research (25-0268).

## Notes

### Competing Interest Statement

The authors have declared no competing interest.

