## Supplementary Data for "A generative network model reveals hierarchical structural memory guiding post-mitotic 3D genome inheritance"

### Extended Data Figures

**Extended Data Fig. 1: HiCONA reconstruction of hierarchical chromatin organization across polymer simulations with varying degrees of compartmentalization and loop-extrusion strength.**

Simulated chromatin contact maps and corresponding hierarchical chromatin organization inferred by HiCONA across combinations of homotypic strength (HS) and loop-extruder release probability (RP). Columns correspond to increasing HS (HS = 0, 1, 2 and 3), controlling the interaction strength between chromatin regions belonging to the same compartment, whereas rows correspond to decreasing RP (RP = 1, 0.5, 0.05, 0.005 and 0.0005), controlling loop-extruder residence time. For each parameter combination, the simulated contact map is shown at the top (log2 counts), followed by the five levels of nested hierarchical chromatin organization inferred by HiCONA (levels 1–5). Colors distinguish individual clusters independently at each hierarchical level.

E

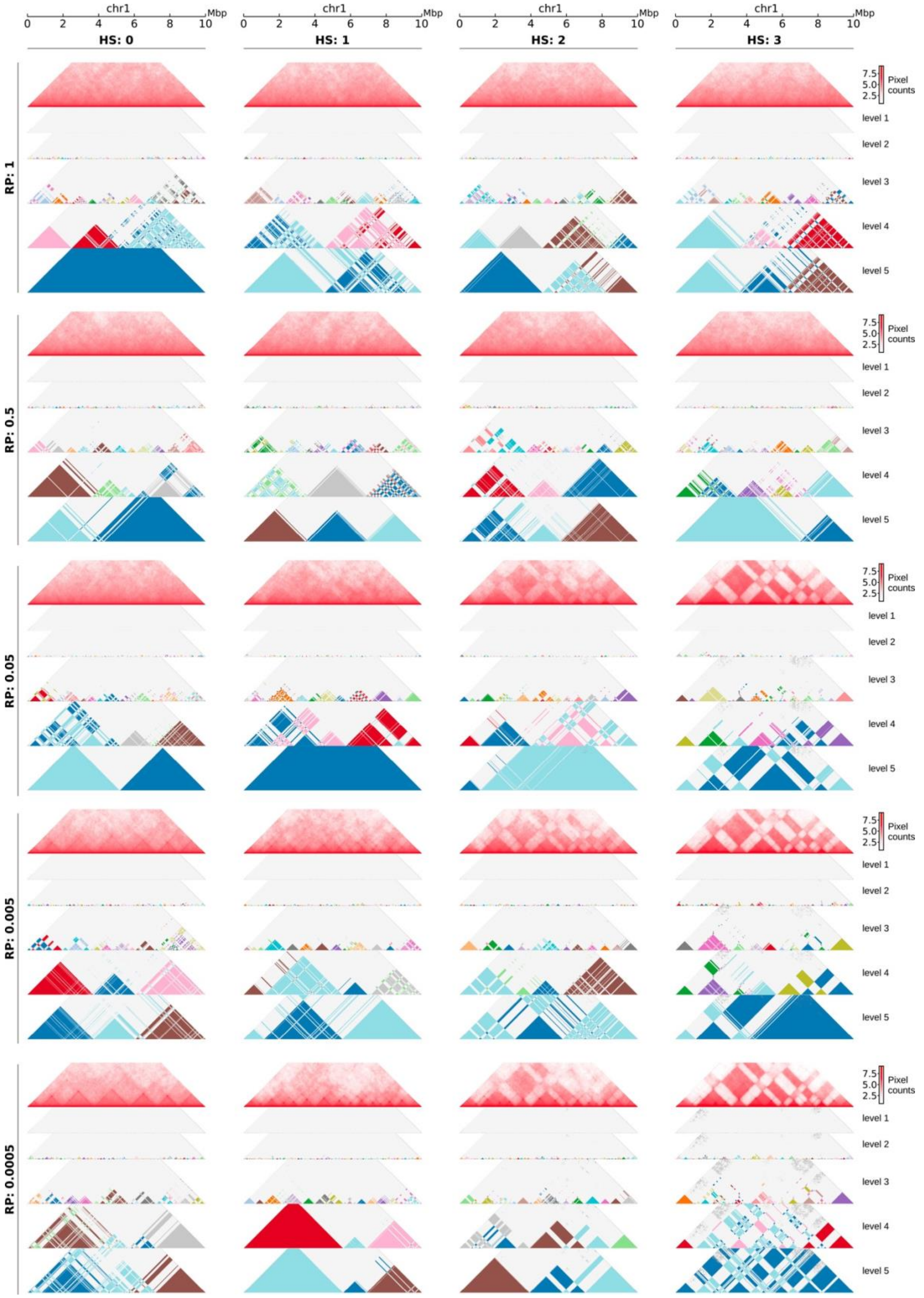
chr1
Mbp
0 2 4 6 8 10
HS: 0
HS: 1
HS: 2
HS: 3
RP: 1
RP: 0.5
RP: 0.05
RP: 0.005
RP: 0.0005
7.5
5.0
2.5
Pixel counts
level 1
level 2
level 3
level 4
level 5

**Extended Data Fig. 2: Extended benchmarking and validation of HiCONA hierarchical chromatin organization.**

**a,** Simulated chromatin contact maps and corresponding hierarchical chromatin organization inferred by HiCONA on a region representing 30Mb of chromatin. **b,** Adjusted Rand index (ARI) between cluster boundaries identified at each hierarchical level of the 30Mb polymer model. **c,** Gini index measuring compartmental homogeneity of clusters at each hierarchical level of the 30Mb polymer model. **d,** Comprehensive benchmarking of HiCONA against 39 TAD-calling methods using TADshop. Heatmap shows the performance of each method across metrics assessing structural protein enrichment, histone modification enrichment, chromatin contact enrichment, and insulation score, together with the overall benchmarking score. Black boxes indicate the best-performing method for each metric. **e,** Chromatin contact map (log2 counts) and nested hierarchical chromatin organization of level1 inferred by HiCONA for chromosome 6 of GM12878 cells at 10-kb resolution used for TAD identification benchmarking. **f,** Chromatin contact map (log2 counts) and nested hierarchical chromatin organization inferred by HiCONA for chromosome 21 of GM12878 cells at 50-kb resolution. **g,** Chromatin contact map (log2 counts) and nested hierarchical chromatin organization inferred by HiCONA for chromosome 13 of GM12878 cells at 50-kb resolution. **h,** Principal component analysis (PCA) of clusters across hierarchical levels of chromosome 13 based on their enrichment across Repli-seq cell-cycle fractions. **i,** Mean Repli-seq z-score profiles of early- and late-replicating clusters identified at each hierarchical level of chromosome 13 across G1b, S1–S4 and G2 cell-cycle fractions. **j,** PCA of clusters across hierarchical levels of chromosome 21 based on their enrichment across Repli-seq cell-cycle fractions. **k,** Mean Repli-seq z-score profiles of early- and late-replicating clusters identified at each hierarchical level of chromosome 21 across G1b, S1–S4 and G2 cell-cycle fractions. **l,** Gini index quantifying compartmental homogeneity at each level of the HiCONA hierarchy for chromosomes 13, 21 and 6 of GM12878 cells.

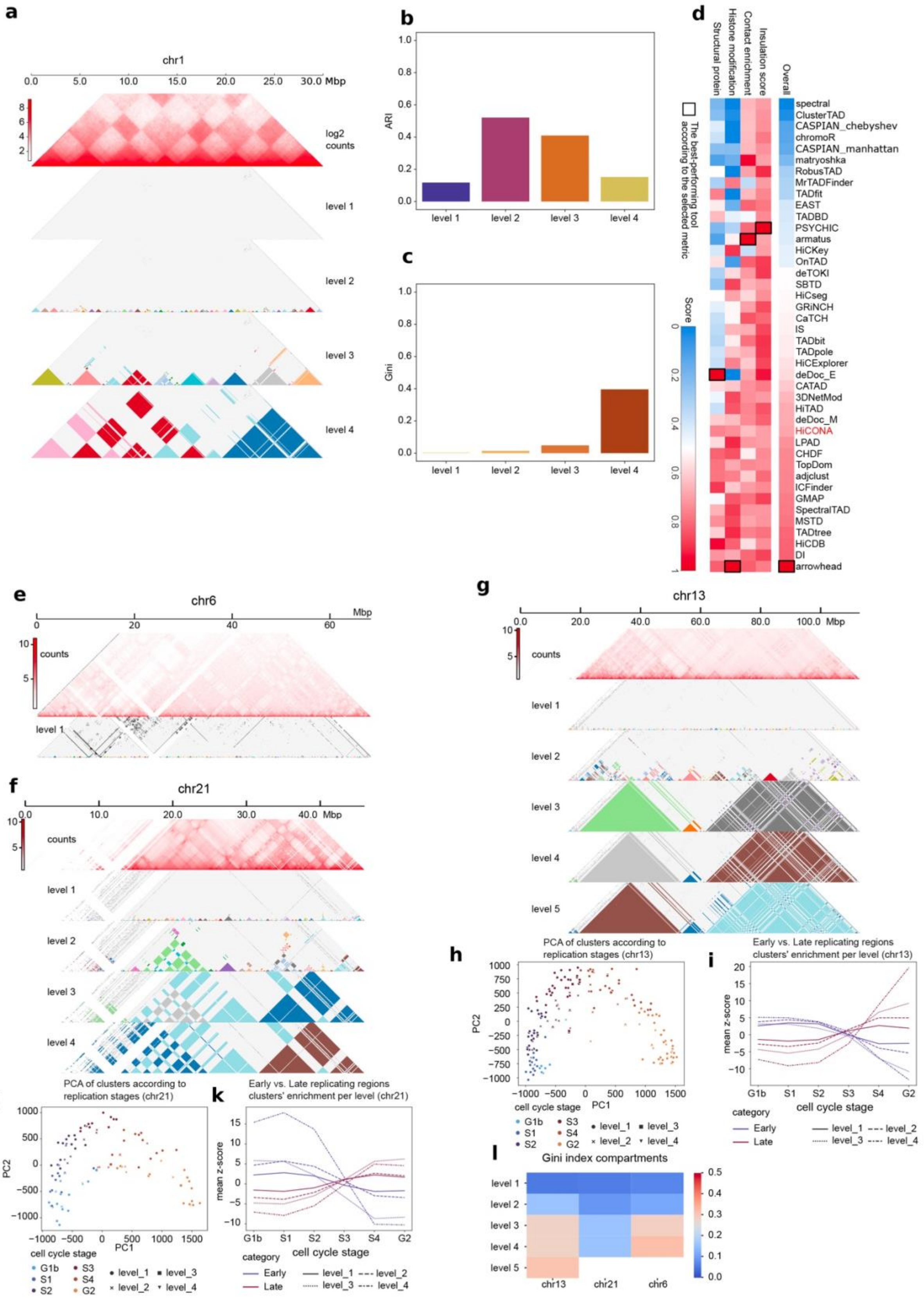
a
chr1
0.0 5.0 10.0 15.0 20.0 25.0 30.0 Mbp
log2 counts
level 1
level 2
level 3
level 4
b
ARI
level 1 level 2 level 3 level 4
c
Gini
level 1 level 2 level 3 level 4
d
Structural protein
Histone modification
Contact enrichment
Insulation score
Overall
The best-performing tool according to the selected metric
Score
spectral
ClusterTAD
CASPIAN_chebyshev
chromoR
CASPIAN_manhattan
matryoshka
RobusTAD
MrTADFinder
TADfit
EAST
TADBD
PSYCHIC
armatus
HiCKey
OnTAD
deTOKI
SBTD
HiCseg
GRiNCH
CaTCH
IS
TADbit
TADpole
HiCExplorer
deDoc_E
CATAD
3DNetMod
HiTAD
deDoc_M
HiCONA
LPAD
CHDF
TopDom
adjclust
ICFinder
GMAP
SpectralTAD
MSTD
TADtree
HiCDB
DI
arrowhead
e
chr6
0 20 40 60 Mbp
counts
level 1
f
chr21
0.0 10.0 20.0 30.0 40.0 Mbp
counts
level 1
level 2
level 3
level 4
g
chr13
0.0 20.0 40.0 60.0 80.0 100.0 Mbp
counts
level 1
level 2
level 3
level 4
level 5
h
PCA of clusters according to replication stages (chr13)
PC1
PC2
cell cycle stage
G1b S1 S2 S3 S4 G2
level_1 level_2 level_3 level_4
i
Early vs. Late replicating regions clusters' enrichment per level (chr13)
mean z-score
cell cycle stage
G1b S1 S2 S3 S4 G2
category
Early
Late
level_1 level_2 level_3 level_4
j
PCA of clusters according to replication stages (chr21)
PC1
PC2
cell cycle stage
G1b S1 S2 S3 S4 G2
level_1 level_2 level_3 level_4
k
Early vs. Late replicating regions clusters' enrichment per level (chr21)
mean z-score
cell cycle stage
G1b S1 S2 S3 S4 G2
category
Early
Late
level_1 level_2 level_3 level_4
l
Gini index compartments
level 1
level 2
level 3
level 4
level 5
chr13 chr21 chr6

**Extended Data Fig. 3: Entropy Impact Score captures compartmental organization across cell types and genomic resolutions.**

**a,b,** Chromatin contact maps (top, log2 counts) and corresponding Entropy Impact Score (EIS) maps (bottom) for chromosome 21 of GM12878 cells at 100-kb (**a**) and 250-kb (**b**) resolution. **c,** Distribution of EIS values for homotypic and heterotypic compartment interactions on chromosome 21 of GM12878 cells at 100-kb and 250-kb resolution. **d–f,** Chromatin contact maps (top, log2 counts) and corresponding EIS maps (bottom) for chromosome 21 of IMR90 cells at 50-kb (**d**), 100-kb (**e**) and 250-kb (**f**) resolution. **g,** Distribution of EIS values for homotypic and heterotypic compartment interactions on chromosome 21 of IMR90 cells at 50-kb, 100-kb and 250-kb resolution. **h,** Two-dimensional density plot showing the relationship between genomic distance and chromatin contact frequency (log2-transformed counts) for chromosome 21 of GM12878 cells at 50-kb resolution. Higher densities are indicated by warmer colors.

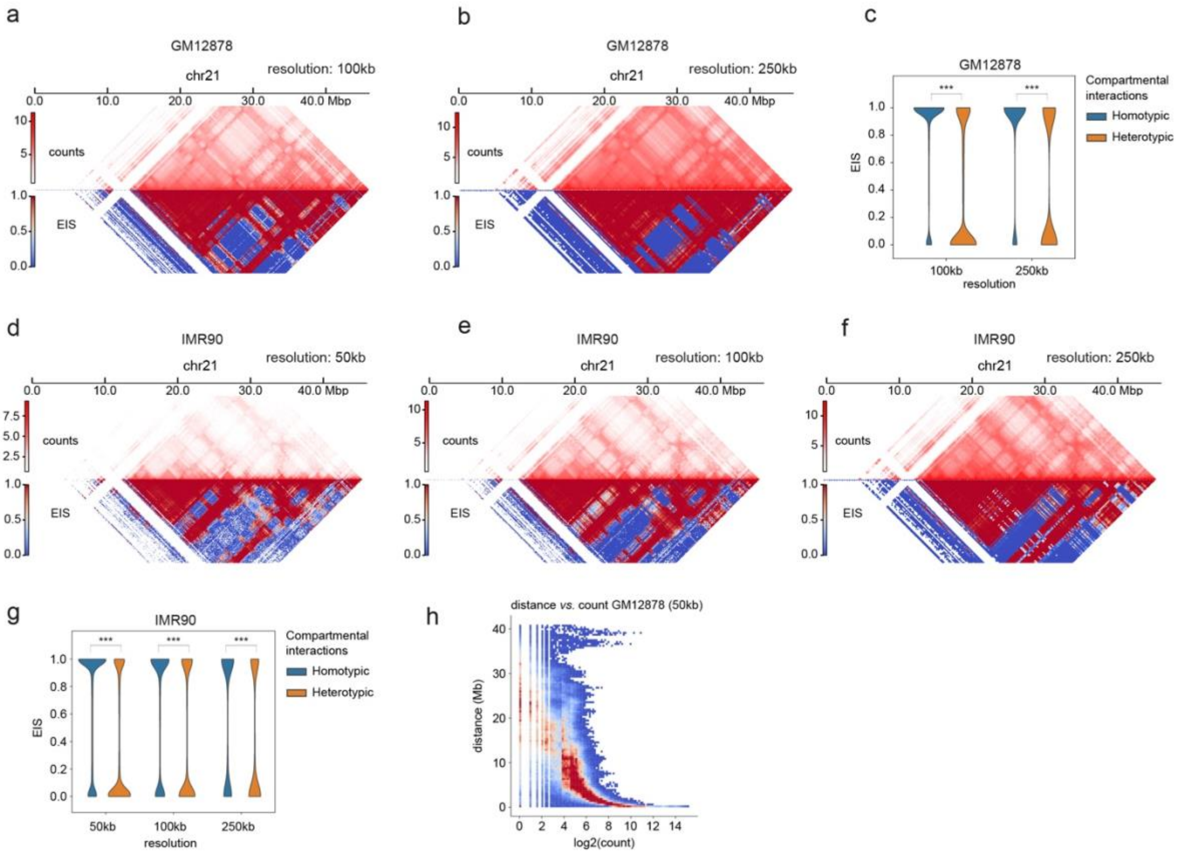
a
GM12878
chr21
resolution: 100kb
0.0 10.0 20.0 30.0 40.0 Mbp
counts
EIS
b
GM12878
chr21
resolution: 250kb
c
GM12878
Compartmental interactions
Homotypic
Heterotypic
EIS
100kb
250kb
resolution
d
IMR90
chr21
resolution: 50kb
e
IMR90
chr21
resolution: 100kb
f
IMR90
chr21
resolution: 250kb
g
IMR90
50kb
100kb
250kb
resolution
h
distance vs. count GM12878 (50kb)
distance (Mb)
log2(count)

**Extended Data Fig. 4: HiCONA captures cohesin-dependent changes in hierarchical chromatin organization and contact impact across genomic scales.**

**a,b,** Chromatin contact maps (log2 counts) and nested hierarchical chromatin organization inferred by HiCONA for chr3:18,000,000–49,000,000 in untreated (**a**) and IAA-treated (**b**) mESCs at 40-kb resolution. The contact map is shown at the top, followed by the hierarchical levels identified by HiCONA. **c,d,** Chromatin contact maps (log2 counts) and corresponding HiCONA hierarchical organization for the Sox2 locus (chr3:33,390,000–36,180,000) in untreated (**c**) and IAA-treated (**d**) mESCs at 20-kb resolution. **e–g,** Chromatin contact maps (top, log2 counts) and corresponding Entropy Impact Score (EIS) maps (bottom) for chr6:31,000,000–61,000,000 in untreated (**e**) and IAA-treated (**f**) mESCs at 40-kb resolution, together with differential maps comparing untreated and IAA-treated conditions (**g**). **h–j,** Chromatin contact maps (top, log2 counts) and corresponding EIS maps (bottom) for chr6:50,600,000–52,800,000 in untreated (**h**) and IAA-treated (**i**) mESCs at 20-kb resolution, together with differential maps comparing untreated and IAA-treated conditions (**j**).

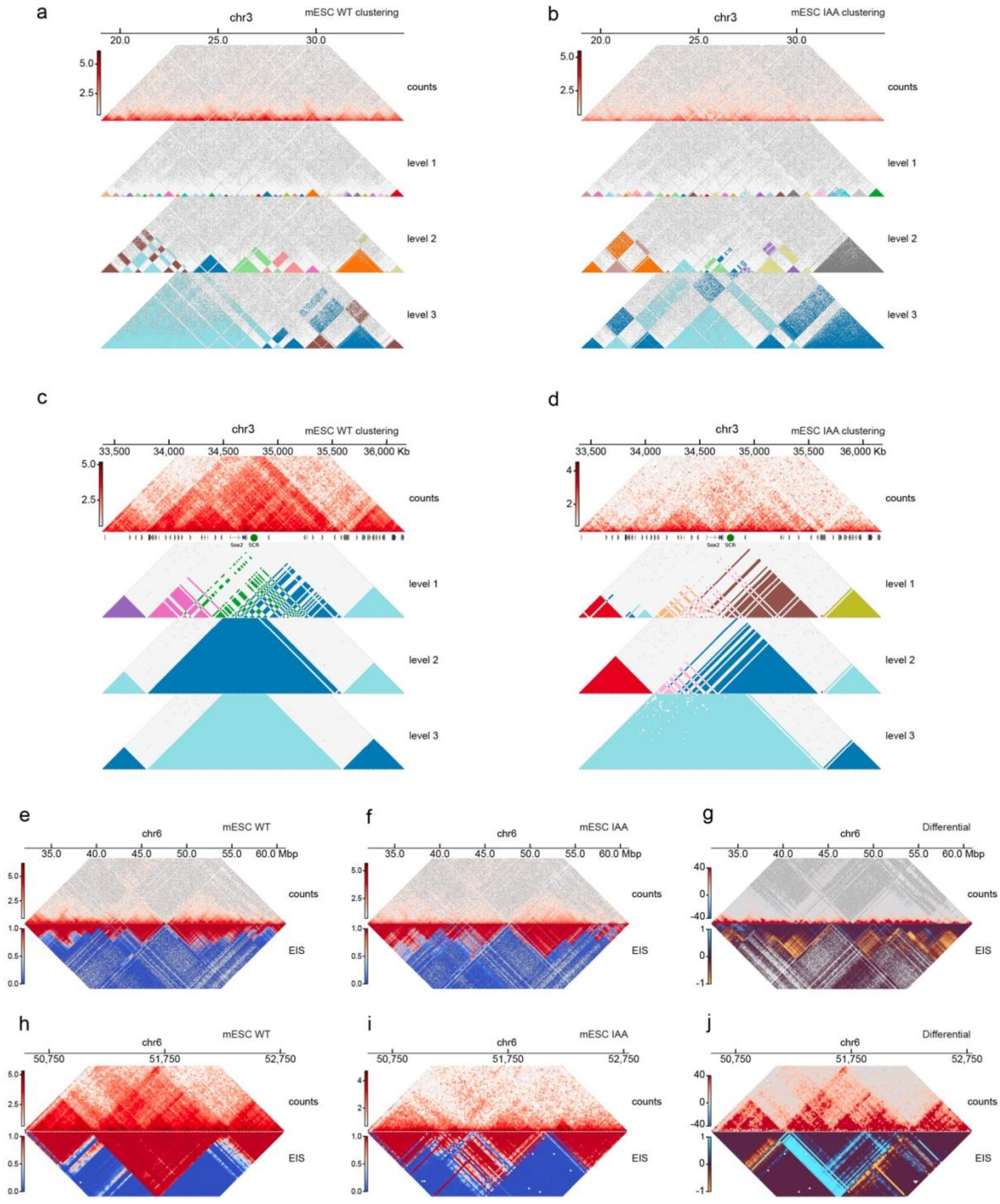
a
chr3
mESC WT clustering
20.0
25.0
30.0
5.0
2.5
counts
level 1
level 2
level 3
b
chr3
mESC IAA clustering
20.0
25.0
30.0
5.0
2.5
counts
level 1
level 2
level 3
c
chr3
mESC WT clustering
33,500
34,000
34,500
35,000
35,500
36,000 Kb
5.0
2.5
counts
Sox2
SCR
level 1
level 2
level 3
d
chr3
mESC IAA clustering
33,500
34,000
34,500
35,000
35,500
36,000 Kb
4
2
counts
Sox2
SCR
level 1
level 2
level 3
e
chr6
mESC WT
35.0
40.0
45.0
50.0
55.0
60.0 Mbp
5.0
2.5
counts
1.0
0.5
0.0
EIS
f
chr6
mESC IAA
35.0
40.0
45.0
50.0
55.0
60.0 Mbp
5.0
2.5
counts
1.0
0.5
0.0
EIS
g
chr6
Differential
35.0
40.0
45.0
50.0
55.0
60.0 Mbp
40
0
-40
counts
1
0
-1
EIS
h
chr6
mESC WT
50,750
51,750
52,750
5.0
2.5
counts
1.0
0.5
0.0
EIS
i
chr6
mESC IAA
50,750
51,750
52,750
4
2
counts
1.0
0.5
0.0
EIS
j
chr6
Differential
50,750
51,750
52,750
40
0
-40
counts
1
0
-1
EIS

**Extended Data Fig. 5: Entropy Impact Score identifies high-impact microcompartment interactions in an independent genomic region.**

**a**, Differential HiC contact map (top, log2 counts) and corresponding ORCA contact map (bottom) comparing untreated and IAA-treated conditions in ORCA dataset. CUT&RUN tracks show enrichment in H3K27ac, H3K4me1, H3K4me3, and H3K27me3. **b–d,** Chromatin contact maps (top, log2 counts) and corresponding Entropy Impact Score (EIS) maps (bottom) for the Klf8 (chr8) region in mESCs under untreated (**b**), transcription inhibition (**c**) and cohesin depletion (**d**) conditions. **e,** Distribution of EIS values for microcompartment interactions compared with all chromatin contacts in WT, TI and IAA-treated mESCs.

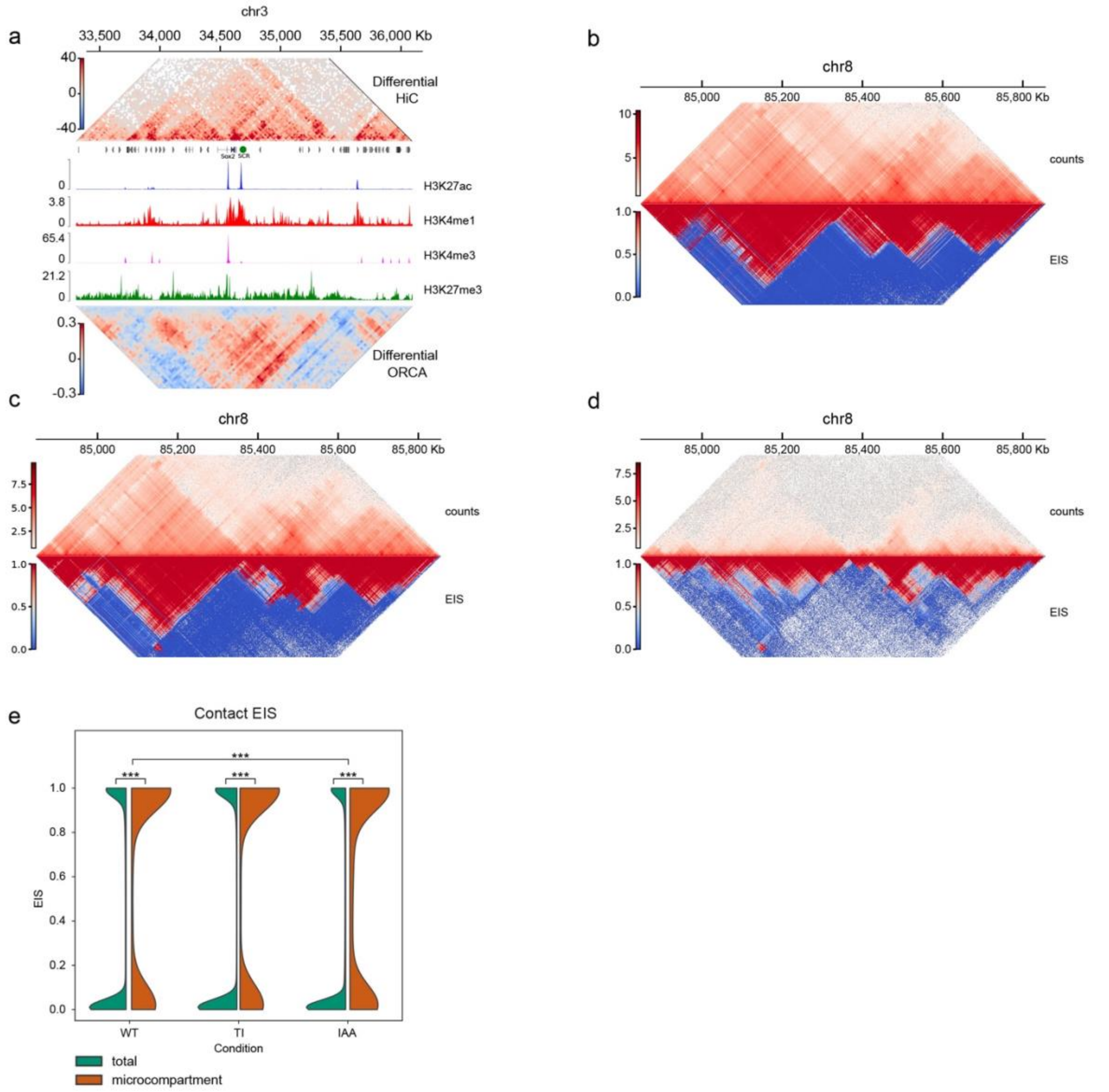


**Extended Data Fig. 6: Dynamic Entropy Impact Score patterns during mitotic exit across additional RCMC loci.**

Chromatin contact maps (top, log2 counts) and corresponding Entropy Impact Score (EIS) maps (bottom) across three additional genomic regions profiled by RCMC during mitotic exit in G1E-ER4 erythroblasts. Columns correspond to chr2:152.0–153.0 Mb (left), chr8:85.0–85.8 Mb (center) and chr8:123.0–124.0 Mb (right). Rows correspond to successive cell-cycle stages: prometaphase (**a–c**), anaphase–telophase (**d–f**), early G1 (**g–i**), intermediate G1 (**j–l**) and late G1 (**m–o**). **p**. Median EIS trajectories throughout cell division for the 5 clusters identified by k-means clustering of microcompartments in G1E-ER4 erythroblasts. **q**. Distribution of chromatin interaction distances within microcompartment clusters. **r**. Log2 fold-change enrichment of asynchronously bound transcription factors (GATA2 and TAL1), histone modifications (H3K36me3, H3K4me1, H3K4me3, H3K27me3, and H3K9me3), RAD21, and CTCF across the cell cycle in clusters identified using only microcompartment contacts.

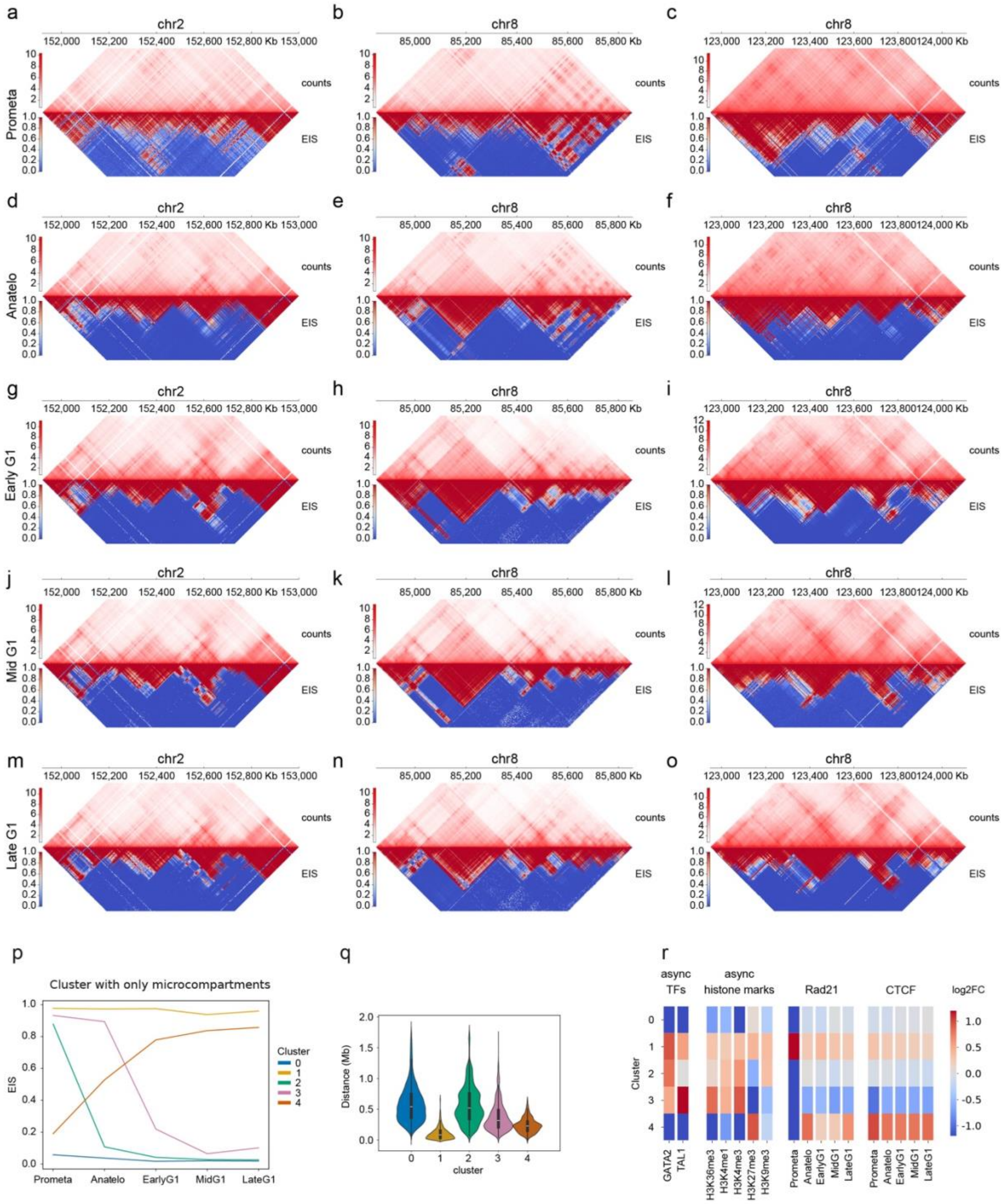

a
b
c
d
e
f
g
h
i
j
k
l
m
n
o
p
q
r
chr2
chr8
152,000 152,200 152,400 152,600 152,800 Kb 153,000
85,000 85,200 85,400 85,600 85,800 Kb
123,000 123,200 123,400 123,600 123,800124,000 Kb
counts
EIS
Prometa
Anatelo
Early G1
Mid G1
Late G1
Cluster with only microcompartments
EIS
Cluster
0
1
2
3
4
Prometa Anatelo EarlyG1 MidG1 LateG1
Distance (Mb)
cluster
async
TFs
async
histone marks
Rad21
CTCF
log2FC
GATA2
TAL1
H3K36me3
H3K4me1
H3K4me3
H3K27me3
H3K9me3
Prometa
Anatelo
EarlyG1
MidG1
LateG1

**Extended Data Fig. 7: Dynamic Entropy Impact Score patterns across additional genomic regions during mitotic exit in DLD-1 cells.**

Chromatin contact maps (top, log2 counts) and corresponding Entropy Impact Score (EIS) maps (bottom) across additional genomic regions in wild-type DLD-1 cells during mitotic exit. Rows correspond to distinct genomic regions: chr6:125–145.5 Mb, chr7:18.5–38.5 Mb, chr7:78–105 Mb, chr8:29.5–59.5 Mb, chr8:100–120 Mb, chr14:33.5–55.5 Mb, chr14:54–80 Mb and chr17:32–52 Mb. Columns correspond to successive cell-cycle stages: prometaphase, anaphase–telophase, cytokinesis, and G1 cells collected 5 h and 10 h after mitotic exit.

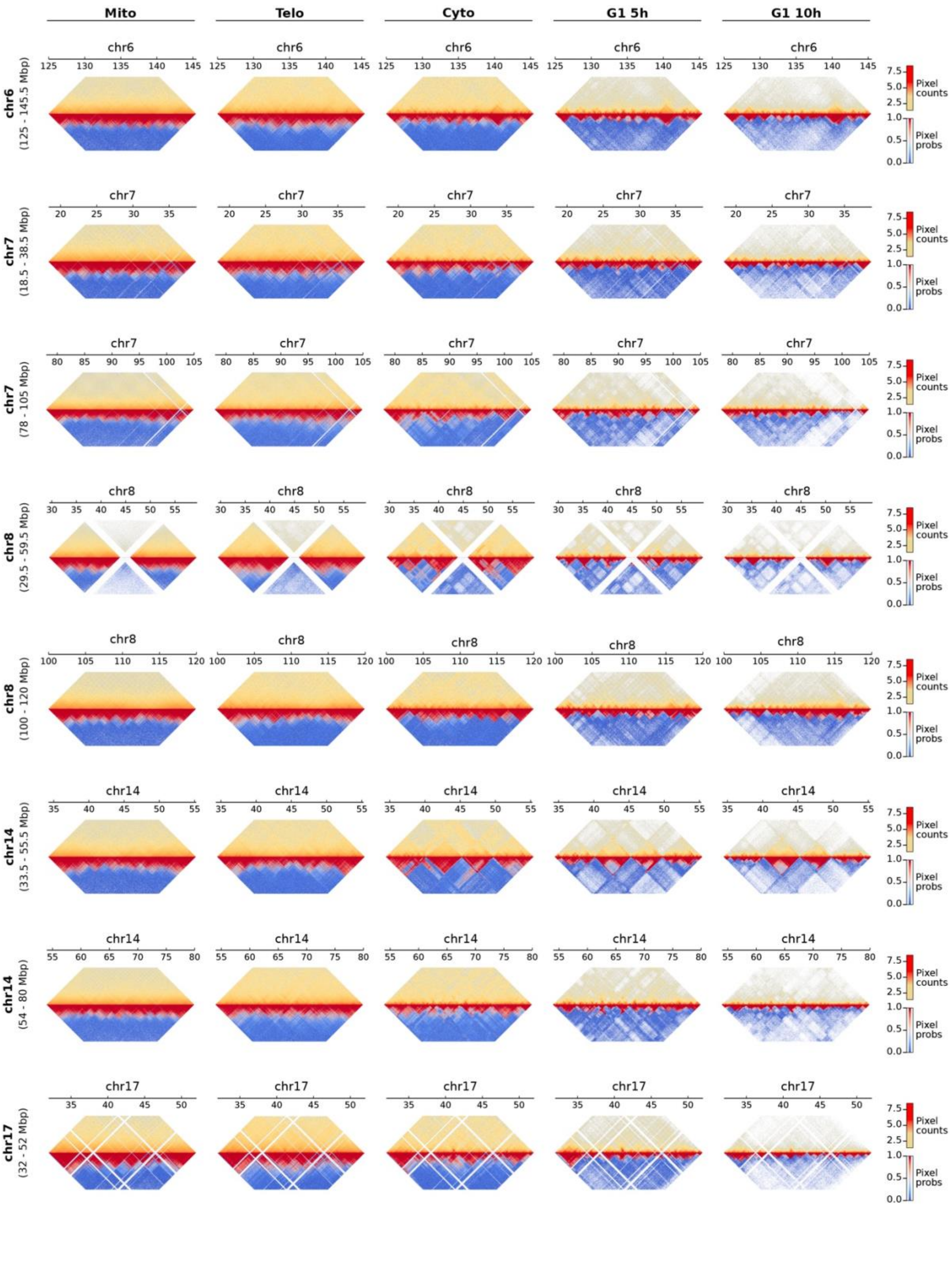

Mito
Telo
Cyto
G1 5h
G1 10h
chr6 (125 - 145.5 Mbp)
chr7 (18.5 - 38.5 Mbp)
chr7 (78 - 105 Mbp)
chr8 (29.5 - 59.5 Mbp)
chr8 (100 - 120 Mbp)
chr14 (33.5 - 55.5 Mbp)
chr14 (54 - 80 Mbp)
chr17 (32 - 52 Mbp)
Pixel counts
Pixel probs

**Extended Data Fig. 8: Relationship between Entropy Impact Score and genomic distance across mitotic exit in DLD-1 cells.**

Two-dimensional density plots showing the relationship between Entropy Impact Score (EIS) and genomic distance of chromatin interactions across multiple genomic regions in wild-type DLD-1 cells. Rows correspond to chr6:125–145.5 Mb, chr7:18.5–38.5 Mb, chr7:78–105 Mb, chr8:29.5–59.5 Mb, chr8:100–120 Mb, chr14:33.5–55.5 Mb, chr14:54–80 Mb, chr17:32–52 Mb and chr17:47.5–73.4 Mb. Columns correspond to successive cell-cycle stages: prometaphase, anaphase–telophase, cytokinesis, and G1 cells collected 5 h and 10 h after mitotic exit. Higher densities are indicated by warmer colors.

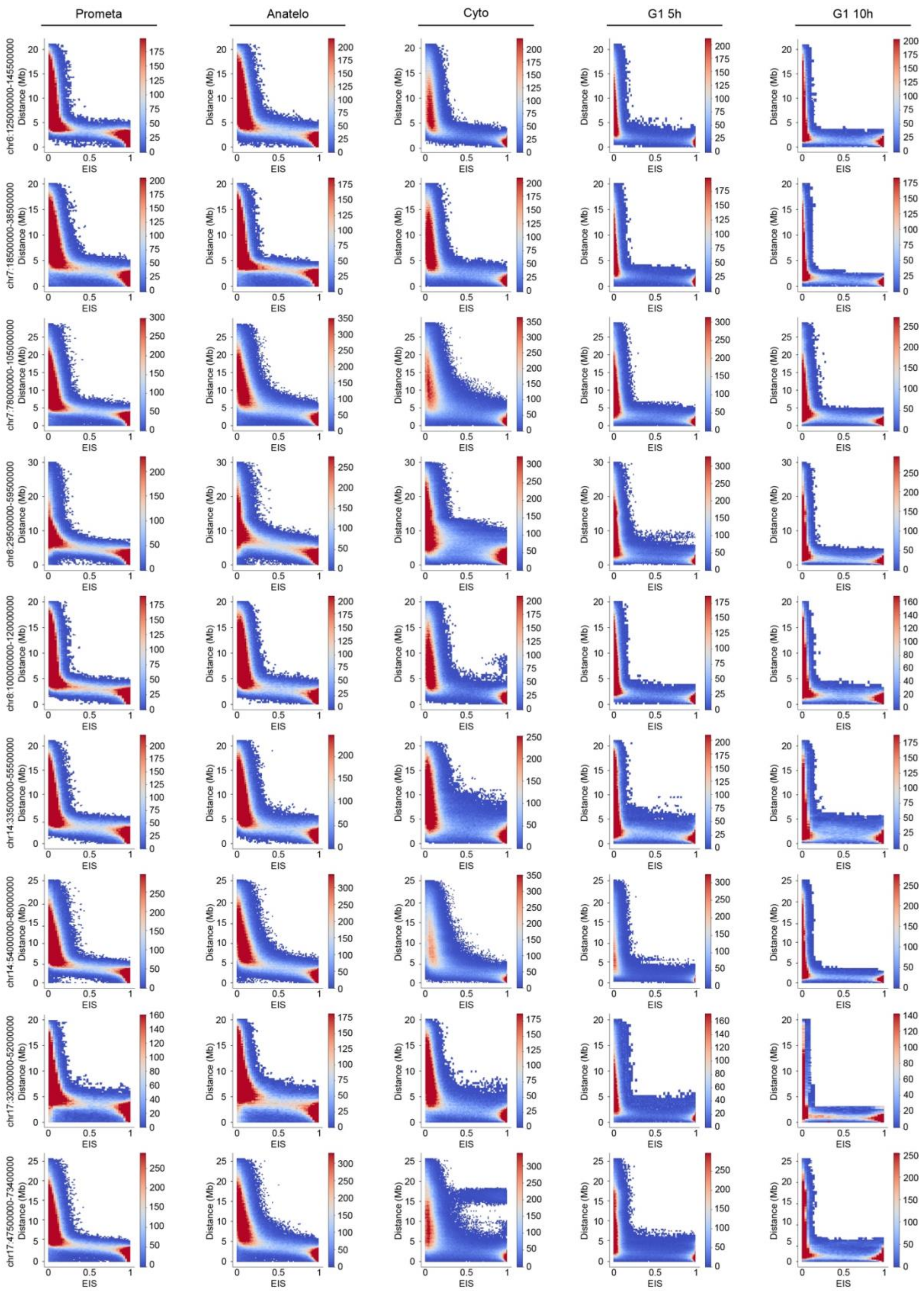
Prometa
Anatelo
Cyto
G1 5h
G1 10h
chr6:125000000-145500000
chr7:18500000-38500000
chr7:78000000-105000000
chr8:29500000-59500000
chr8:100000000-120000000
chr14:33500000-55500000
chr14:54000000-80000000
chr17:32000000-52000000
chr17:47500000-73400000
Distance (Mb)
EIS

**Extended Data Fig. 9: Genomic distribution of EIS-defined chromatin interaction classes in DLD-1 cells.**

**a–i,** Genomic distribution of chromatin contacts classified according to their Entropy Impact Score (EIS) trajectories throughout cell-cycle progression and their subcompartmental association in untreated (top) and deltaGAP1 (bottom) DLD-1 cells. Contacts are shown for chr6:125–145.5 Mb (**a**), chr7:18.5–38.5 Mb (**b**), chr7:78–105 Mb (**c**), chr8:29.5–59.5 Mb (**d**), chr8:100–120 Mb (**e**), chr14:33.5–55.5 Mb (**f**), chr14:54–80 Mb (**g**), chr17:32–52 Mb (**h**) and chr17:47.5–73.4 Mb (**i**). **j,** Color key indicating contact classes according to subcompartment pairing (AA, BB or AB) and genomic interaction range (short, intermediate or long).

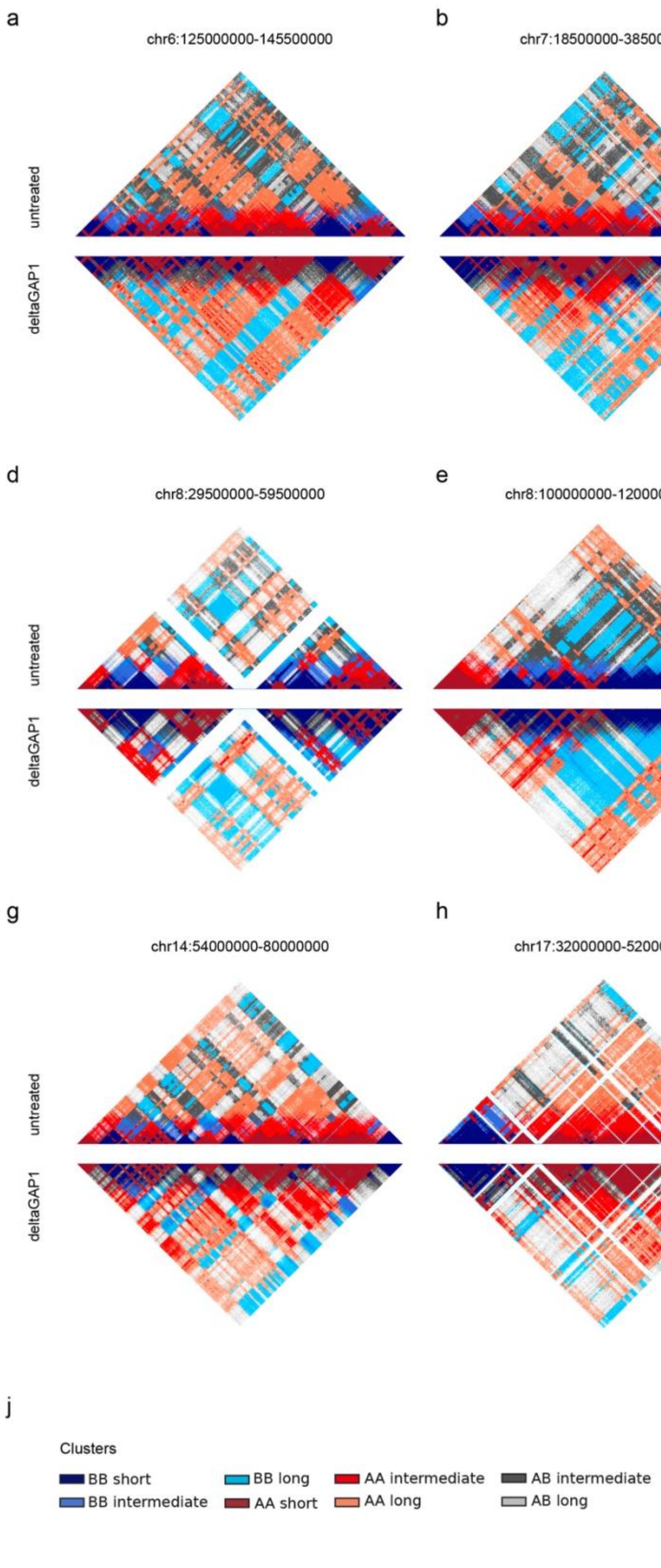
a
chr6:125000000-145500000
b
chr7:18500000-38500000
c
chr7:78000000-105000000
untreated
deltaGAP1
d
chr8:29500000-59500000
e
chr8:100000000-120000000
f
chr14:33500000-55500000
g
chr14:54000000-80000000
h
chr17:32000000-52000000
i
chr17:47500000-73400000
j
Clusters
BB short
BB intermediate
BB long
AA short
AA intermediate
AA long
AB intermediate
AB long

**Extended Data Fig. 10: Cell-cycle dynamics of chromatin interactions associated with histone modifications and CTCF in DLD-1 cells.**

**a–d,** Aggregate peak analysis (APA) of chromatin contacts associated with H3K4me3 (**a**), H3K27ac (**b**), H3K27me3 (**c**) and CTCF (**d**) across mitotic exit in wild-type DLD-1 cells. Contacts are stratified according to the EIS-defined chromatin interaction clusters and according to ChIP–seq peaks identified in G1 or prometaphase. For each cluster, observed/expected APA maps are shown across prometaphase, anaphase–telophase, cytokinesis, G1 5 h, and G1 10 h. **e,** Saddle plots of subcompartment interaction enrichment for the clusters identified in deltaGAP1 cells.

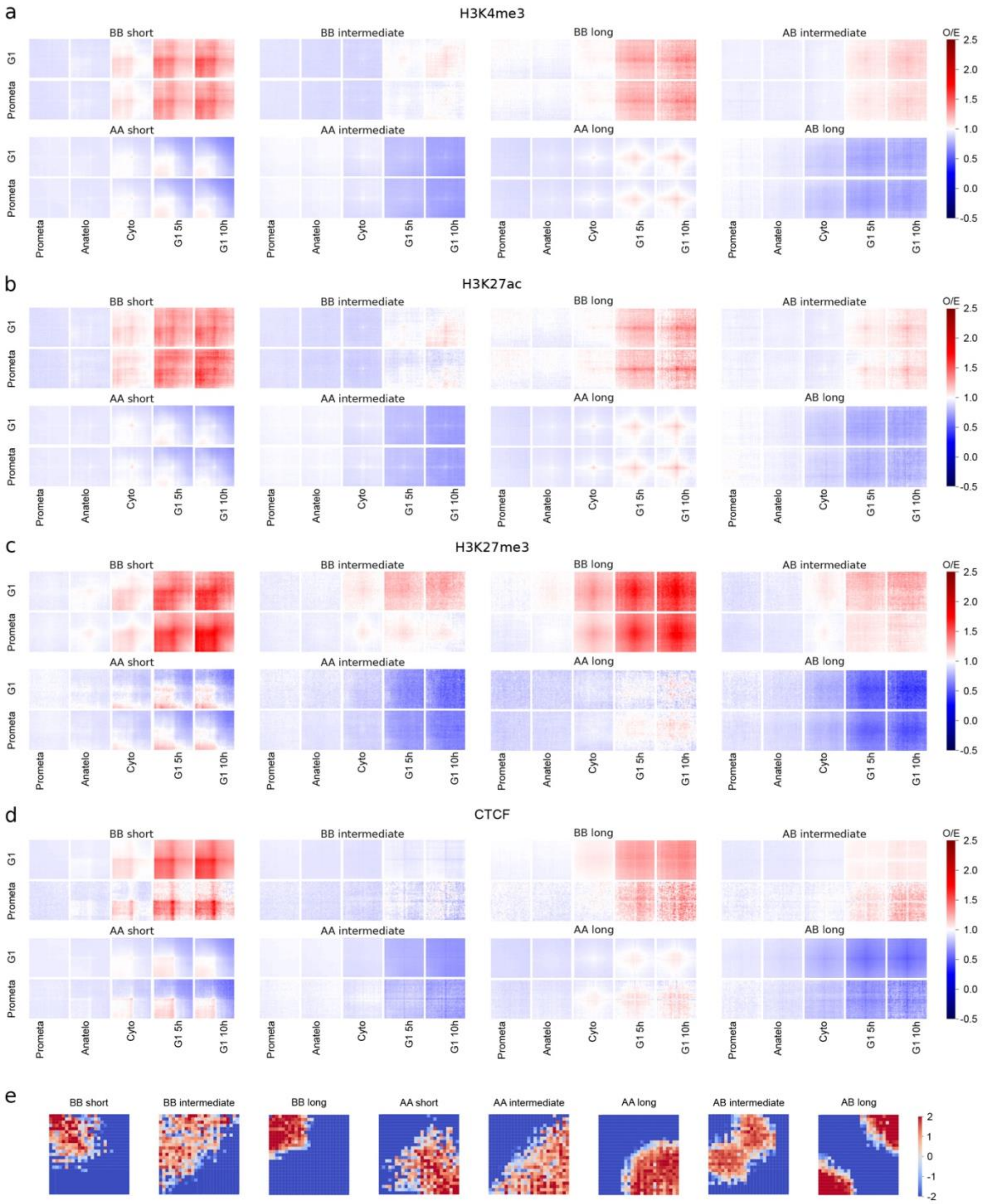

a
H3K4me3
BB short
BB intermediate
BB long
AB intermediate
AA short
AA intermediate
AA long
AB long
G1
Prometa
Prometa
Anatelo
Cyto
G1 5h
G1 10h
O/E
2.5
2.0
1.5
1.0
0.5
0.0
-0.5
b
H3K27ac
c
H3K27me3
d
CTCF
e
BB short
BB intermediate
BB long
AA short
AA intermediate
AA long
AB intermediate
AB long
2
1
0
-1
-2

**Extended Data Fig. 11: Dynamic Entropy Impact Score patterns across additional genomic regions during mitotic exit in deltaGAP1 DLD-1 cells.**

Chromatin contact maps (top, log2 counts) and corresponding Entropy Impact Score (EIS) maps (bottom) across additional genomic regions in deltaGAP1 DLD-1 cells during mitotic exit. Rows correspond to distinct genomic regions: chr6:125–145.5 Mb, chr7:18.5–38.5 Mb, chr7:78–105 Mb, chr8:29.5–59.5 Mb, chr8:100–120 Mb, chr14:33.5–55.5 Mb, chr14:54–80 Mb and chr17:32–52 Mb. Columns correspond to successive cell-cycle stages: prometaphase, anaphase–telophase, cytokinesis, and G1 cells collected 5 h and 10 h after mitotic exit.

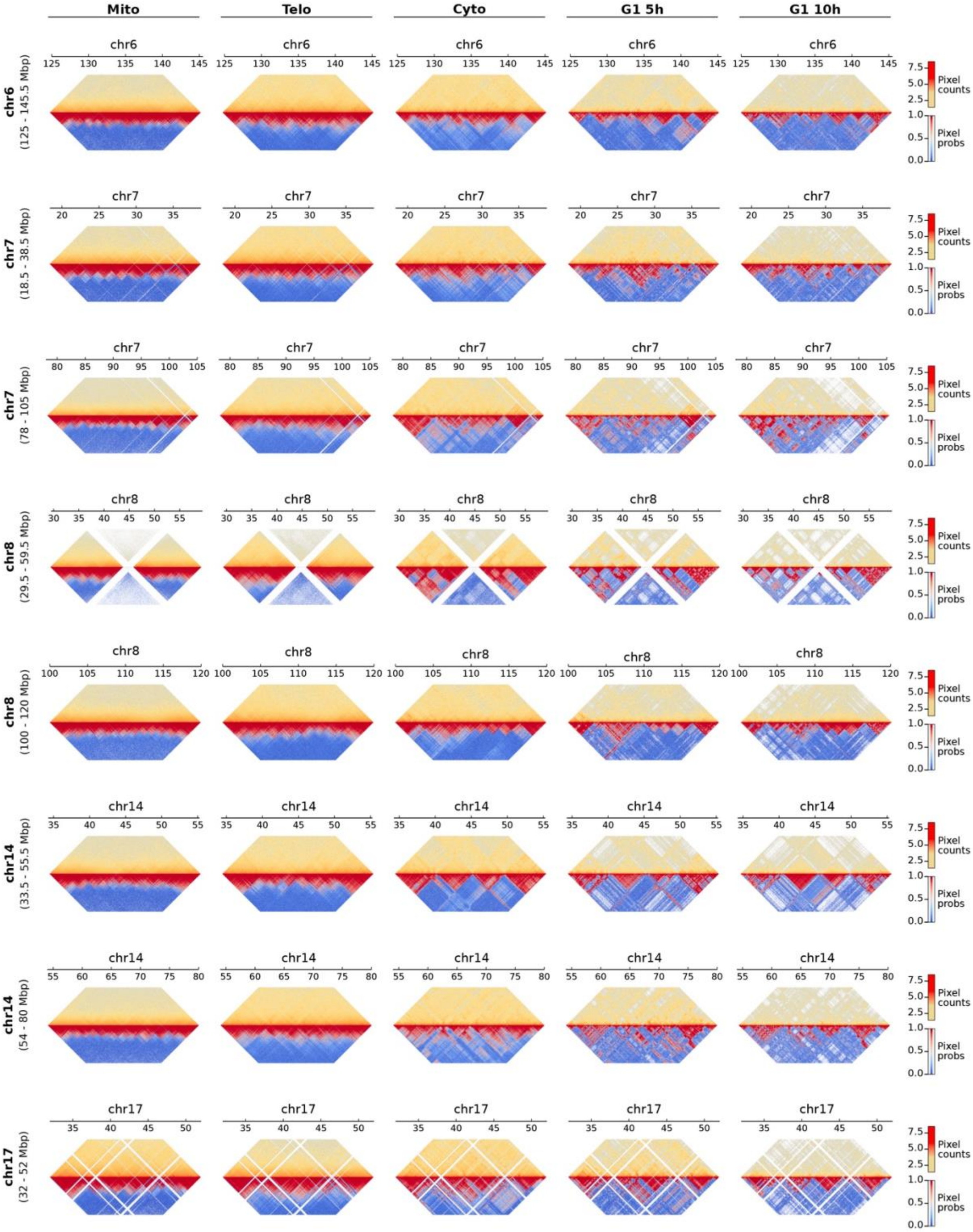

Mito
Telo
Cyto
G1 5h
G1 10h
chr6 (125 - 145.5 Mbp)
chr7 (18.5 - 38.5 Mbp)
chr7 (78 - 105 Mbp)
chr8 (29.5 - 59.5 Mbp)
chr8 (100 - 120 Mbp)
chr14 (33.5 - 55.5 Mbp)
chr14 (54 - 80 Mbp)
chr17 (32 - 52 Mbp)
Pixel counts
Pixel probs

**Extended Data Fig. 12: Relationship between Entropy Impact Score and genomic distance across mitotic exit in deltaGAP1 DLD-1 cells.**

Two-dimensional density plots showing the relationship between Entropy Impact Score (EIS) and genomic distance of chromatin interactions across multiple genomic regions in deltaGAP1 DLD-1 cells. Rows correspond to chr6:125–145.5 Mb, chr7:18.5–38.5 Mb, chr7:78–105 Mb, chr8:29.5–59.5 Mb, chr8:100–120 Mb, chr14:33.5–55.5 Mb, chr14:54–80 Mb, chr17:32–52 Mb and chr17:47.5–73.4 Mb. Columns correspond to successive cell-cycle stages: prometaphase, anaphase–telophase, cytokinesis, and G1 cells collected 5 h and 10 h after mitotic exit. Warmer colors indicate higher densities.

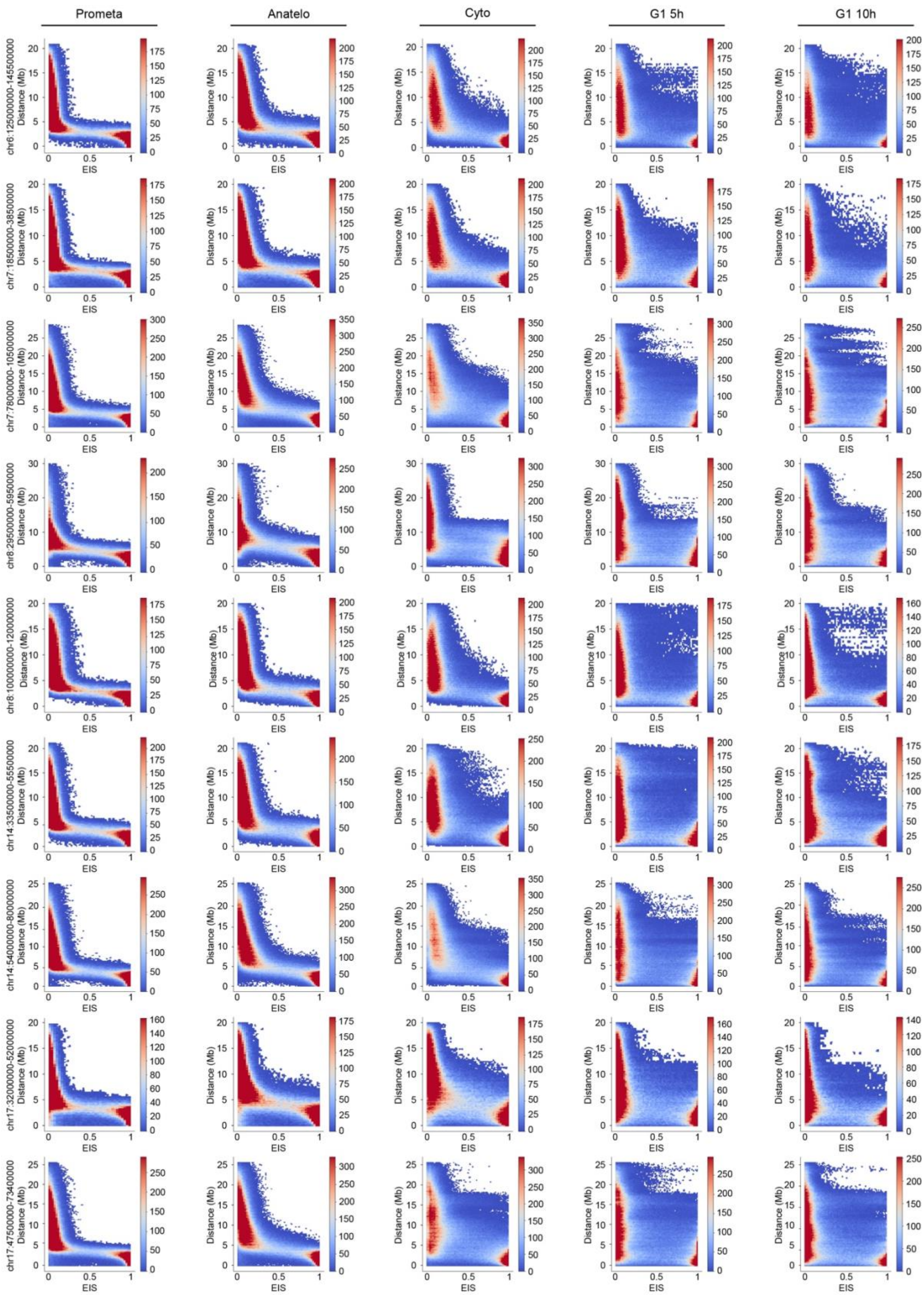

Prometa
Anatelo
Cyto
G1 5h
G1 10h
chr6:125000000-145500000
chr7:18500000-38500000
chr7:78000000-105000000
chr8:29500000-59500000
chr8:100000000-120000000
chr14:33500000-55500000
chr14:54000000-80000000
chr17:32000000-52000000
chr17:47500000-73400000
Distance (Mb)
EIS

**Extended Data Fig. 13: Chromatin interaction dynamics of additional deltaGAP1-defined clusters and nascent RNA-associated contacts during mitotic exit.**

**a–c,** Aggregate peak analysis (APA) of chromatin contacts across mitotic exit in untreated and deltaGAP1 DLD-1 cells for the EIS-defined clusters. APA maps are shown for contacts associated with mitotic nascent RNA (**a**), H3K27ac (**b**) and H3K27me3 (**c**) across prometaphase, anaphase–telophase, cytokinesis, G1 5 h and G1 10 h. **d,** APA-score trajectories across mitotic exit for contacts associated with both mitotic nascent RNA and prometaphase H3K27ac peaks, stratified by EIS-defined cluster. **e,** APA-score trajectories for contacts associated with both mitotic nascent RNA and prometaphase H3K4me3 peaks. **f,** APA-score trajectories for contacts associated with both mitotic nascent RNA and G1 H3K27ac peaks. **g,** APA-score trajectories for mitotic nascent RNA-associated contacts lacking prometaphase H3K27ac peaks. In **d–g**, trajectories are shown for all EIS-defined chromatin interaction clusters. **h**, Venn diagram showing the overlap of significant GO terms between Huh7, *Δ*GAP1 and untreated DLD-1 cells. **i**, Lolliplot representing GO terms clustered according to GO slim classes divided in shared, DLD-1 specific, and Huh7 specific.

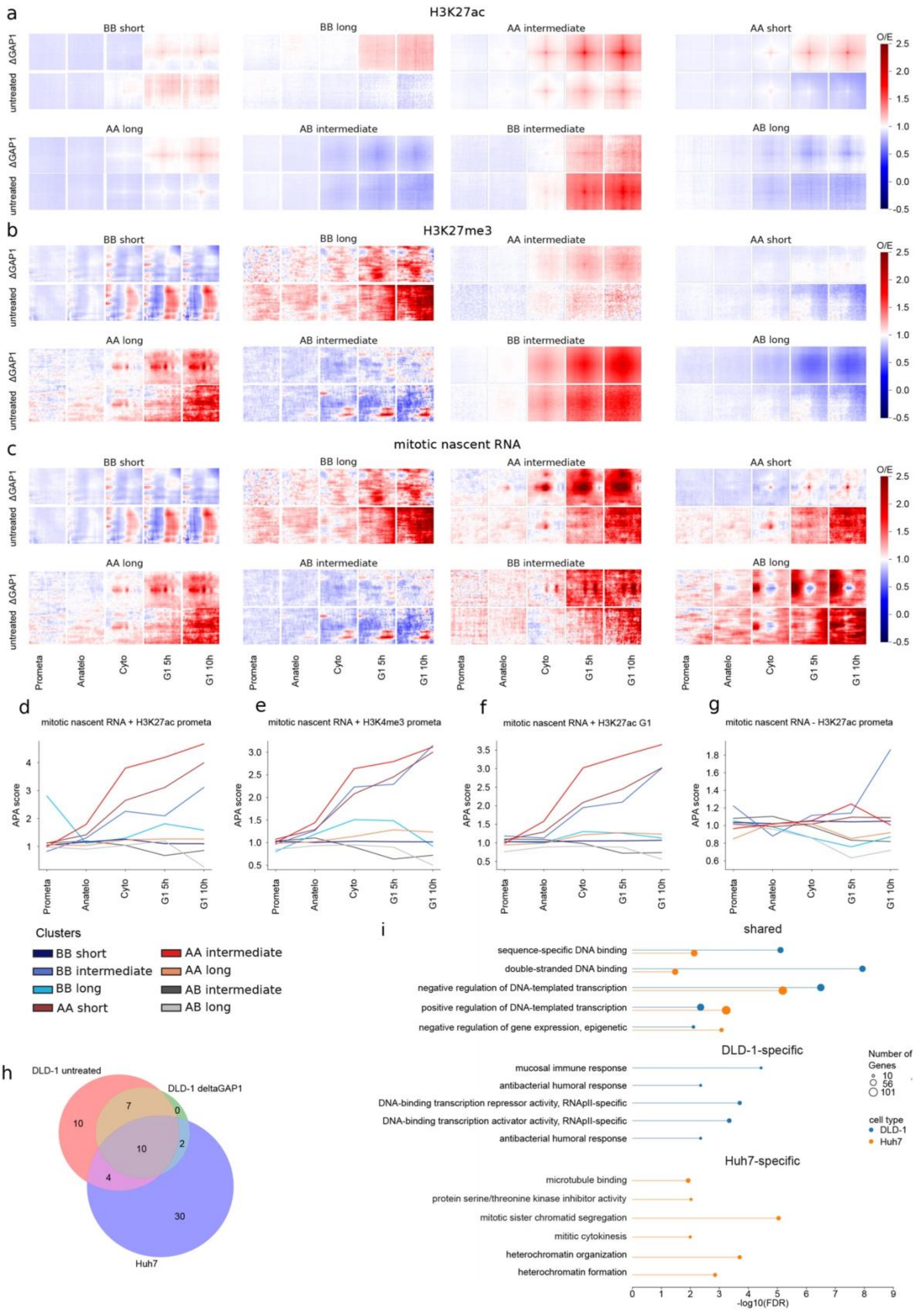
a
H3K27ac
BB short
BB long
AA intermediate
AA short
AA long
AB intermediate
BB intermediate
AB long
ΔGAP1
untreated
O/E
2.5
2.0
1.5
1.0
0.5
0.0
-0.5
b
H3K27me3
c
mitotic nascent RNA
Prometa
Anatelo
Cyto
G1 5h
G1 10h
d
mitotic nascent RNA + H3K27ac prometa
e
mitotic nascent RNA + H3K4me3 prometa
f
mitotic nascent RNA + H3K27ac G1
g
mitotic nascent RNA - H3K27ac prometa
APA score
Clusters
BB short
BB intermediate
BB long
AA short
AA intermediate
AA long
AB intermediate
AB long
h
DLD-1 untreated
DLD-1 deltaGAP1
Huh7
10
7
0
10
2
4
30
i
shared
sequence-specific DNA binding
double-stranded DNA binding
negative regulation of DNA-templated transcription
positive regulation of DNA-templated transcription
negative regulation of gene expression, epigenetic
DLD-1-specific
mucosal immune response
antibacterial humoral response
DNA-binding transcription repressor activity, RNApII-specific
DNA-binding transcription activator activity, RNApII-specific
antibacterial humoral response
Huh7-specific
microtubule binding
protein serine/threonine kinase inhibitor activity
mitotic sister chromatid segregation
mititic cytokinesis
heterochromatin organization
heterochromatin formation
Number of Genes
10
56
101
cell type
DLD-1
Huh7
-log10(FDR)

### Supplementary Tables

#### Supplementary Table 1

List of GO terms significantly represented in mitotic transcripts of untreated DLD-1 cells.

#### Supplementary Table 2

List of GO terms significantly represented in mitotic transcripts of *Δ*DLD-1 cells.

#### Supplementary Table 3

List of GO terms significantly represented in mitotic transcripts of Huh-7 cells.

#### Supplementary Table 4

List of slimGO terms significantly represented in mitotic transcripts of Huh-7 cells vs. DLD-1 cells.

#### Supplementary Table 5

List of chromatin regions analyzed: cell type, ID of the dataset, region analyzed, and contact matrix resolution.

#### Supplementary Table 6

Chromatin network parameters related to HiCONA runtime: number of edges, total EIS, total counts, and chromatin region size.
